# Long-term serial passaging of SARS-CoV-2 reveals signatures of convergent evolution

**DOI:** 10.1101/2023.11.02.565396

**Authors:** Charles S.P. Foster, Gregory J. Walker, Tyra Jean, Maureen Wong, Levent Brassil, Sonia R. Isaacs, Yonghui Lyu, Stuart Turville, Anthony Kelleher, William D. Rawlinson

## Abstract

Understanding viral evolutionary dynamics is crucial to pandemic responses, prediction of virus adaptation over time, and virus surveillance for public health strategies. Whole-genome sequencing (WGS) of SARS-CoV-2 has enabled fine-grained studies of virus evolution in the human population. Serial passaging *in vitro* offers a complementary controlled environment to investigate the emergence and persistence of genetic variants that may confer selective advantage. In this study, nine virus lineages, including four “variants of concern” and three former “variants under investigation”, were sampled over ≥33 serial passages (range 33-100) in Vero E6 cells. WGS was used to examine virus evolutionary dynamics and identify key mutations with implications for fitness and/or transmissibility. Viruses accumulated mutations regularly during serial passaging. Many low-frequency variants were lost but others became fixed, suggesting either *in vitro* benefits, or at least a lack of deleterious effect. Mutations arose convergently both across passage lines, and when compared with contemporaneous SARS-CoV-2 clinical sequences. These mutations included some hypothesised to drive lineage success through host immune evasion (e.g. S:A67V, S:H655Y). The appearance of these mutations *in vitro* suggested key mutations can arise convergently even in the absence of a multicellular host immune response through mechanisms other than immune-driven mutation. Such mutations may provide other benefits to the viruses *in vitro*, or arise stochastically. Our quantitative investigation into SARS-CoV-2 evolutionary dynamics spans the greatest number of serial passages to date, and will inform measures to reduce the effects of SARS-CoV-2 infection on the human population.

**Importance:** The ongoing evolution of SARS-CoV-2 remains a challenge for long term public health efforts to minimise the effects of COVID-19. Whole-genome sequencing of outbreak cases has enabled global contact tracing efforts and the identification of mutations of concern within the virus’ genome. However, complementary approaches are necessary to inform our understanding of virus evolution and clinical outcomes. Here we charted evolution of the virus within a controlled cell culture environment, focusing on nine different virus lineages. Our approach demonstrates how SARS-CoV-2 continues to evolve readily in vitro, with changes mirroring those seen in outbreak cases globally. Findings of the study are important for i) investigating the mechanisms of how mutations arise, ii) predicting the future evolutionary trajectory of SARS-CoV-2, and iii) informing treatment and prevention design.

## Introduction

The COVID-19 pandemic saw whole-genome sequencing (WGS) embraced on an unprecedented scale, with nearly 100 countries worldwide possessing SARS-CoV-2 WGS capability and contributing to publicly available sequencing repositories (1). Consequently, the number of available high quality full-genome sequences for SARS-CoV-2 (>17,000,000 as of April 2025) far exceeds that of any other pathogen. The accessibility of sequencing data has allowed the population-scale evolution of SARS-CoV-2 to be characterised with fine-grained detail. This has facilitated real-time monitoring of the spread of SARS-CoV-2, identification of new viral lineages, and provided crucial insights into virus evolution. For example, the rise and fall of different SARS-CoV-2 lineages has been charted, with attempts to link spread of virus strains with the presence of key mutations associated with lineage success (2).

Global sequencing efforts have revealed that SARS-CoV-2 has continued to accumulate mutations since first emergence in late 2019. The rate of mutation accumulation within SARS-CoV-2 (∼1D×D10^−6^ to 2D×D10^−6^Dmutations per nucleotide per replication cycle) is typical of betacoronaviruses. However, this is below the rate typical in other RNA viruses that lack proofreading mechanisms (2). Retrospective consideration can divide the COVID-19 pandemic into several ‘eras’: the initial phase of the pandemic characterised by apparently limited evolution of the virus, the sudden emergence of the highly divergent virus lineages with altered phenotypes (variants of concern, VOCs) (3), and periods of gradual evolution within VOC lineages (3, 4). Determining the evolutionary forces responsible for each of these eras is key to understanding how the pandemic has progressed over time, how it will continue to progress as SARS-CoV-2 shifts towards being an endemic virus (5), and how future pandemics might unfold.

Analysis of the changes that have occurred in SARS-CoV-2 genomes from the global clinical population suggests the predominant driver of evolution in SARS-CoV-2 has been the accumulation of mutations resulting in increased transmissibility (Markov et al., 2023). Increased transmissibility is a multifaceted trait resulting from a given virus having an increased ability to survive within an infected individual, shed from an infected individual, establish within a new individual, or some combination of each of these abilities. These are all influenced by virus evading the host immune response. A range of ‘key mutations’ that are partially and additively responsible for each of these components of transmissibility have been identified. For example, several mutations in the Spike region have been linked to enhanced receptor binding, such as D614G, identified early in the pandemic (6), and N501Y, which was identified in several VOCs (7–9).

Whilst genomic data derived from routine genomic sequencing for clinical or surveillance purposes (henceforth referred to as the “clinical population” of SARS-CoV-2) has enhanced understanding the evolution of SARS-CoV-2, there are inherent limitations in relying solely on this information. The availability of WGS data is influenced by i) differential sequencing efforts between countries, with few data from low- and middle-income countries (1), ii) when data sharing in online repositories is not consistent among jurisdictions, iii) multiple uncontrolled processes such as natural selection, genetic drift, host immunity, and population dynamics, and iv) virus dependent factors (persistence, replication competence). It is challenging to understand the adaptation of SARS-CoV-2 to specific selective pressures, or to predict the potential evolutionary pathways, when relying solely on the clinical population. Serial passaging *in vitro* provides complementary insights into the ongoing evolution of viruses. In these experiments, the virus is passaged through successive generations within a cell line, allowing study of virus evolution in the absence of immune and therapeutic pressures, as mutations accumulate over time during the course of passaging (10). This process is often used to develop virus stocks, assess therapeutics (11), assess virus attenuation for vaccine development (12), or study the evolutionary trajectory of viruses over long term experiments, within strictly controlled time periods (13, 14).

Few studies have investigated the evolutionary dynamics of SARS-CoV-2 over time *in vitro*. Based on limited passaging of ancestral lineages (<15 passages), it was reported that SARS-CoV-2 accumulates mutations readily *in vitro*, especially when passaged in Vero E6 cells, with an estimated *in vitro* spontaneous mutation rate of 1.3 × 10^−6^ ± 0.2 × 10^−6^ per-base per-infection cycle (15). These studies generally have studied a relatively small number of passages (e.g. <10), over a short time, and relatively few SARS-CoV-2 lineages. The greatest number of lineages published to date is 15 passages with two lineages from early in the pandemic (15). There is evidence for the convergent acquisition of mutations among samples taken through serial passage, including some that have been observed in the global clinical population of SARS-CoV-2 (16). For example, a common observation is the accumulation of mutations associated with Vero E6-cultured samples (15), especially in the furin cleavage site (PRRAR) within the S1/S2 domain of the SARS-CoV-2 spike glycoprotein. The furin cleavage site has been proposed as a key component of the pathogenesis of SARS-CoV-2 (17), with an important role in the affinity of SARS-CoV-2 for human hosts (18). Deletion of the furin cleavage site has been linked with a fitness advantage in Vero E6 cells, albeit with this advantage reduced when ectopic expression of TMPRSS2 is introduced (18). Consequently, SARS-CoV-2 propagated in Vero E6 cell lines has been observed to repeatedly develop mutation(s) in the furin cleavage site, likely as an adaptation to the loss of the cell-surface serine protease-mediated cell entry mode since Vero E6 cells do not express TMPRSS2 (19). Accordingly, unintended selective pressures to develop alternative modes of cell entry can be imparted on Vero E6-propagated SARS-CoV-2, emphasising the need to interpret results in the context of known cell line-specific impacts.

Further serial passaging studies are needed to assess the evolutionary patterns of SARS-CoV-2 *in vitro* with those in the human population, including the *de novo* appearance and/or fixation of mutations. In this study, we compared the accumulation of mutations *in vitro* in 11 different passaged viruses, corresponding to nine unique Pango lineages, throughout long-term serial passaging (33-100 passages per virus line). The evolutionary dynamics of these passage lines was determined by characterising the mutations that accumulated, both in terms of their number and potential functional consequences. Additionally, we assessed mutations for evidence of convergence across passage lines and/or with key mutations from the global clinical population of SARS-CoV-2.

## Methods

### Rationale and study design

Since the beginning of the COVID-19 pandemic, routine whole-genome sequencing (WGS) has been conducted at the Virology Research Laboratory, Serology and Virology Division (SAViD), NSW Health Pathology East, on PCR-positive SARS-CoV-2 clinical specimens. Viruses from clinical specimens were isolated *in vitro*, and then serially passaged and sequenced to investigate the evolutionary dynamics of SARS-CoV-2 in culture (see additional methods below). Initially an early variant (A.2.2) was chosen for this purpose. Later, additional lineages were also grown *in vitro* based on their relevance within the clinical population globally, with a particular focus on ‘variants of concern’ (VOC) or ‘variants under investigation’ (VUI), as designated by the World Health Organisation. Eleven passage lines were established and were taken through a minimum of 33 and up to a maximum of 100 passages (Table 1).

**Table 1:**
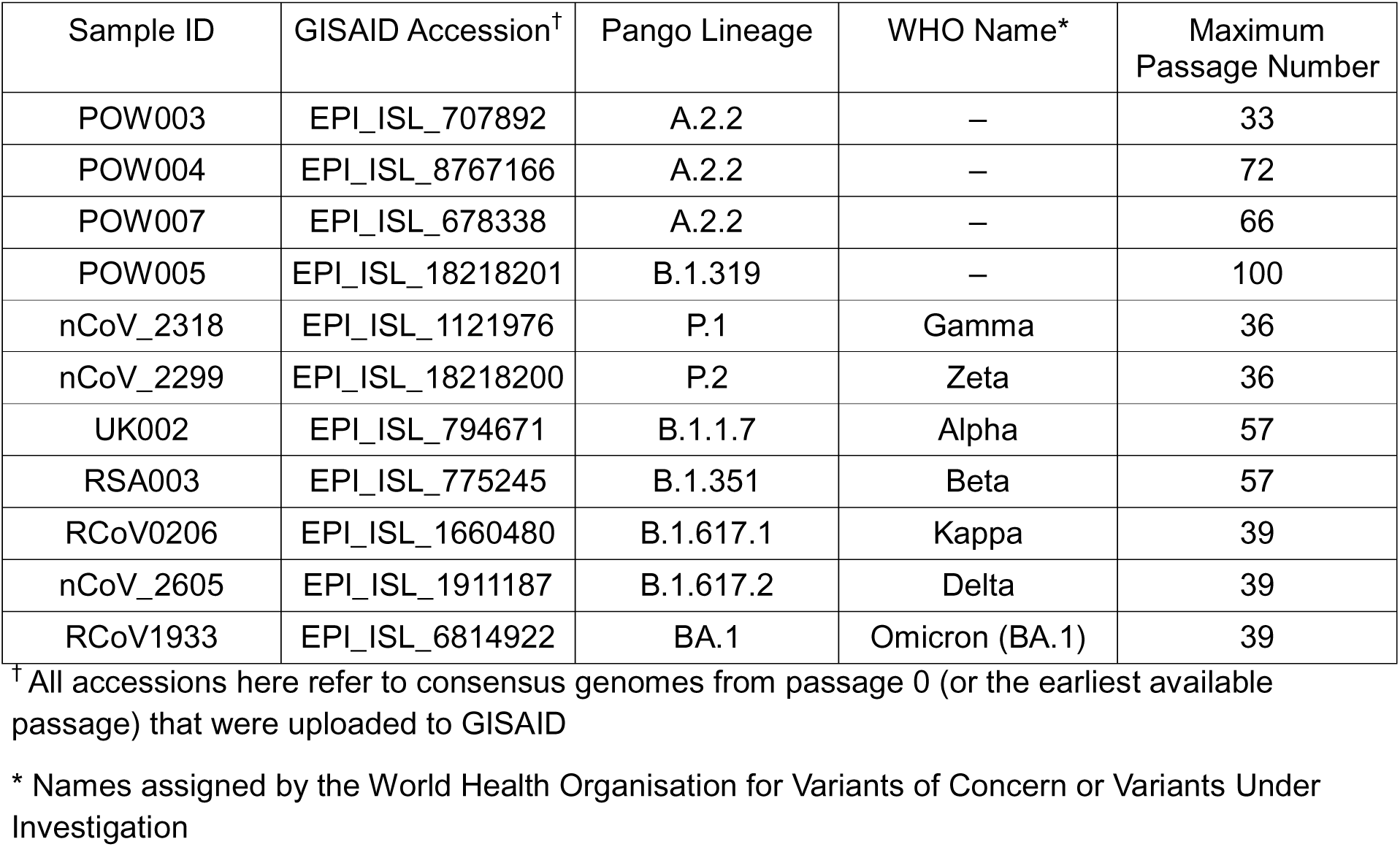
Overview of the passage lines used within this study, including their nomenclatural assignments and the number of serial passages.

| Sample ID | GISAID Accession<sup>†</sup> | Pango Lineage | WHO Name* | Maximum Passage Number |
| --- | --- | --- | --- | --- |
| POW003 | EPI_ISL_707892 | A.2.2 | – | 33 |
| POW004 | EPI_ISL_8767166 | A.2.2 | – | 72 |
| POW007 | EPI_ISL_678338 | A.2.2 | – | 66 |
| POW005 | EPI_ISL_18218201 | B.1.319 | – | 100 |
| nCoV_2318 | EPI_ISL_1121976 | P.1 | Gamma | 36 |
| nCoV_2299 | EPI_ISL_18218200 | P.2 | Zeta | 36 |
| UK002 | EPI_ISL_794671 | B.1.1.7 | Alpha | 57 |
| RSA003 | EPI_ISL_775245 | B.1.351 | Beta | 57 |
| RCoV0206 | EPI_ISL_1660480 | B.1.617.1 | Kappa | 39 |
| nCoV_2605 | EPI_ISL_1911187 | B.1.617.2 | Delta | 39 |
| RCoV1933 | EPI_ISL_6814922 | BA.1 | Omicron (BA.1) | 39 |

<sup>†</sup> All accessions here refer to consensus genomes from passage 0 (or the earliest available passage) that were uploaded to GISAID
\* Names assigned by the World Health Organisation for Variants of Concern or Variants Under Investigation

### Serial passaging

All SARS-CoV-2 cultures were performed in Vero E6 cells that were purchased commercially and have been verified by the ECACC (ECACC catalogue number 85020206). Vero E6 cells were maintained (up to a maximum of passage 30) in Minimum Essential Media supplemented with 10%-FBS and 1xPSG (MEM-10), and incubated at 37°C, 5% CO_2_. The day prior to infection, a 24-well plate was seeded with 120,000 cells per well and incubated overnight, for approximately 90% confluency at the time of virus inoculation. The Vero E6 cell line was chosen as it supports viral replication to high titres due to its lack of interferon production and relatively high expression of the ACE-2 receptor (20). The appearance of any immune evasion-associated mutations from the clinical population during in vitro passaging in Vero E6 cells allowed inference of fitness advantages of these mutations beyond immune evasion.

Clinical specimens (nasopharyngeal swabs in Virus Transport Media, referred to as “Passage 0”) positive using diagnostic RT-qPCR were transported to a PC3 facility for virus culture. Specimens were spin-filtered (0.22 µm sterile Ultrafree-MC Centrifugal Filter) to remove cellular debris and reduce potential for bacterial and fungal contamination. Cell culture maintenance media was removed from wells, and 100 µL of virus-containing flow-through from spin-filtered specimens was added to wells. Plates were incubated for 1hr, before inoculum was removed and replaced with 500 µL maintenance media (MEM-2). Virus cultures were incubated for 3-4 days and observed for cytopathic effect (CPE). Cultured virus was then harvested (passage 1), diluted 1:1000, and used to inoculate a fresh plate of naïve Vero E6 cells (passage 2). This dilution was chosen for an estimated MOI of 0.01, based upon viral titres plateauing in Vero E6 cells at approximately 2×10^6^ TCID50/mL. Cultures of each SARS-CoV-2 lineage were maintained up to a maximum of passage 100. Aliquots of each virus passage were frozen at −80°C until later extraction for whole-genome sequencing.

### Whole-genome sequencing

Samples from each passage line underwent whole-genome sequencing (WGS) as described previously (21), with the exception of the choice of amplicon scheme. In the present study, amplification was performed using the “Midnight” protocol, comprising tiled 1200 nucleotide amplicons (22), followed by 150 bp paired-end sequencing on an Illumina MiSeq (Illumina Inc). Sequences were obtained from passages 0–6, then every three passages onwards where possible, up to a minimum of 33 passages (Table 1).

### Quality control and variant calling

Sequencing data were processed and analysed using an in-house bioinformatics pipeline (“CIS”) (23), which leverages open-source bioinformatics tools for whole-genome sequencing, with workflow management controlled by Snakemake (24). Briefly, initial QC of sequencing reads was performed with fastp v0.23.2 (25), with removal of residual sequencing adapters, error correction, and filtration of reads based on a quality score threshold. Reads passing QC were then mapped against the SARS-CoV-2 reference genome (NC_045512.2) using bwa mem v0.7.17-r1188 (26), and amplicon primer regions were soft-clipped using ivar v1.3.1 (27). Probabilistic realignment of reads and variant calling was performed with lofreq v2.1.5 (28). A consensus genome was assembled using bcftools v1.13 (29), and coverage statistics estimated using bedtools v2.30.0 (30). Finally, Pango lineages were assigned using pangolin (31) and Nextclade (32), then relevant pipeline statistics and additional QC metrics by Nextclade were incorporated into a final report. The pipeline is freely available from https://github.com/charlesfoster/covid-illumina-snakemake.

Several of the default parameters in the CIS pipeline were optimised for the present study. The minimum depth of coverage for a variant to be called was retained at 10, but we reduced minimum SNP and indel variant allele frequency parameters to 0.01 (later filtered further, see below). While this allowed tracking of low-frequency variants, we acknowledge that a lack of uniformity in the depth of coverage across the SARS-CoV-2 genome might impact upon the minimum callable variant allele frequency across sites. Consequently, there is a chance that some variants occurring at very low allele frequencies might be missed in areas of low depth of coverage, but this is a usual limitation when investigating low-frequency variants using whole-genome sequencing in general.

### Tracking mutational changes during serial passaging

We assessed the changes in called variants within and among passage lines throughout serial passaging using an in-house “vartracker” pipeline (see below, available from https://github.com/charlesfoster/vartracker), using the variant call format (VCF) files from lofreq as input. For each passage sample within a passage line, these VCF files were derived by calling variants against the NC_045512.2 SARS-CoV-2 reference genome (using the CIS pipeline, as per above), not against the prior passage number. Doing so allows variants to be discussed using standard notation for SARS-CoV-2, while also allowing the gain and/or loss of variants between passages to be tracked.

The vartracker pipeline comprises an in-house python script to allow interrogation of the mutations that accumulate within a passage line during serial passaging. During the pipeline, the VCF files for each passage line were prefiltered to retain variants with a minimum single-nucleotide variant (SNV) and indel frequency 0.1 and 0.25, respectively, using cyvcf2 (33) and bcftools.

Functional annotation of called variants was inferred using bcftools csq (34), followed by merging into a single file. The merged VCF was then processed using custom python code to track the change in presence/absence of a given variant throughout the trajectory of the passage line (from passage 0 to the final passage), as well as the change in allele frequency of each variant throughout passaging. Variants were classified under four main broad categories: present in passage 0 but lost throughout passaging (‘original_lost’), present in passage 0 and retained throughout passaging (‘original_retained’), absent in passage 0 but gained and retained throughout passaging (‘new_persistent’), or absent in passage 0 but gained and lost throughout passaging (‘new_transient’). All variants were also subject to a suite of quality control measures incorporating the estimated variant depth, the overall site depth, the depth in a window of sites surrounding the variant site, and were assigned an overall metric of quality (PASS/FAIL).

We investigated convergent evolution of any new_persistent or new_transient variants among passage lines. For the purposes of this study, we defined convergence as (a) any case where any two (or more) replicates obtained the same mutation independently during the course of serial passaging, or (b) any passage line developed a mutation recognised as clinically significant within the global population of SARS-CoV-2 based on searches against a database of SARS-CoV-2 mutations comprising information on the functional consequences of mutations (35). Additionally, we calculated the rate of mutation accumulation within each passage line, considering the rate among the overall genome as well as individual genes/regions within the genome, using the following formula: 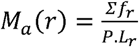 (following the approach of (15)). In this formula, Σ*f_r_* is the sum of the frequency of all detected mutations in a region *r*, *P* is the number of passages, and *L_r_* is the length of *r*. We calculated the rate for each applicable passage up to and including passage 33 to observe whether the inferred mutation rate varies over time.

We investigated the evolutionary dynamics within all passage lines from passage 0, or the earliest passage successfully sequenced, until their final passage number (Table 1). However, for all comparisons among passage lines, we restricted our focus to passages 0–33, with passage 33 representing the maximum passage number in common that all passage lines had reached. We also investigated in further detail the evolutionary trajectory of passage line “POW005” (Pango lineage: B.1.319), which has been maintained for the greatest number of passages (100) across all passage lines. The full results for all other passage lines across all passages are not discussed in this manuscript, but are available within the supplementary materials (Table S2). It is also worth noting that sequencing data for some passage numbers from some passage lines are not available. In these cases, particularly with samples corresponding to VOCs, there was insufficient material from the early passages to conduct WGS and/or the material was needed for unrelated neutralization assay studies.

### Phylogenetic analysis

A phylogenetic tree was estimated to investigate further the evolution of the passaged virus lines *in vitro*. All isolate consensus genomes were aligned against the Wuhan Hu-1 reference genome using minimap2 (36), followed by trimming of the 5’ and 3’ UTRs and conversion to multifasta format using gofasta (37). We then used an in-house script to mask known problematic sites from the alignment (38), and filtered out sites with too many gaps using esl-alimask (hmmer.org). Finally, we inferred a maximum likelihood phylogeny for the samples using iqtree2 (39), with the best fitting nucleotide substitution model chosen using a constrained search using ModelFinder (‘mset’: GTR, TIM2, GTR; ‘mfreq’: F; ‘mrate’: ‘G,R’) (40), and support estimated using 1000 ultrafast bootstraps (41).

## Results and Discussion

### Sequencing metrics

The majority of samples from each passage line were successfully sequenced (Supplementary Table S1). Excluding one sample that yielded no sequence, the mean reference genome coverage at a depth of at least 10 reads was 99.19% (SD: 1.07). Each of these samples was also sequenced at a high mean depth (mean 1334.48, SD: 753). All except one passage line were called as the same lineage as their original isolate (passage 0) after their maximum number of passages. The exception was “POW005”, which was originally called as the lineage B.1.319, but from passage 5 onwards was called as B.1 by pangolin, albeit with conflicting placements by Usher (B.1(1/3); B.1.319(1/3); B.1.616(1/3)). However, Nextclade called the lineage of passage POW005 as B.1.319 from passage 0 through to passage 100, despite the accumulation of many private mutations. The number of sequenced passages varied between passage lines (Table 1), including up to 100 passages for “POW005” (see below). Sequencing metrics and results for all passage numbers for all passage lines are in Supplementary Table S1–2. Here, all comparisons among passage lines are discussed based on a maximum of 33 passages to allow a common point of comparison. The exception is the POW005 passage line, for which the results over 100 passages are discussed.

### Evolutionary dynamics across passage lines *in vitro*

The passaged isolates steadily accumulated mutations *in vitro* throughout the genome with fluctuating variant allele frequencies (VAFs), reflecting clinical isolates (Figure 1, Supplementary Table S2). Many of these mutations were detected at a very low VAF within a given passage (e.g., <10%) and were then lost in subsequent passages. Conversely, some new mutations that arose during passaging reached a consensus-level frequency (≥50% VAF) and/or persisted until the final passage at near-fixation at a consensus-level frequency (Figure 2, Figure 3, Supplementary Table S2. Supplementary Figures S1–S26). Most variants (∼90%) relative to the Wuhan Hu-1 reference genome present in passage 0 of a given passage line were retained during the course of serial passaging. Occasionally these ‘original’ variants were lost, most commonly when they were originally present at a sub-consensus level, but sometimes even when well supported (allele frequency >50%, ∼2% of all variants). Variants were detected in all genes of the SARS-CoV-2 genome but occurred at different frequencies among genes and among passage lines (Figures 1– 4, Supplementary Figures S1–26).

**Figure 1:**
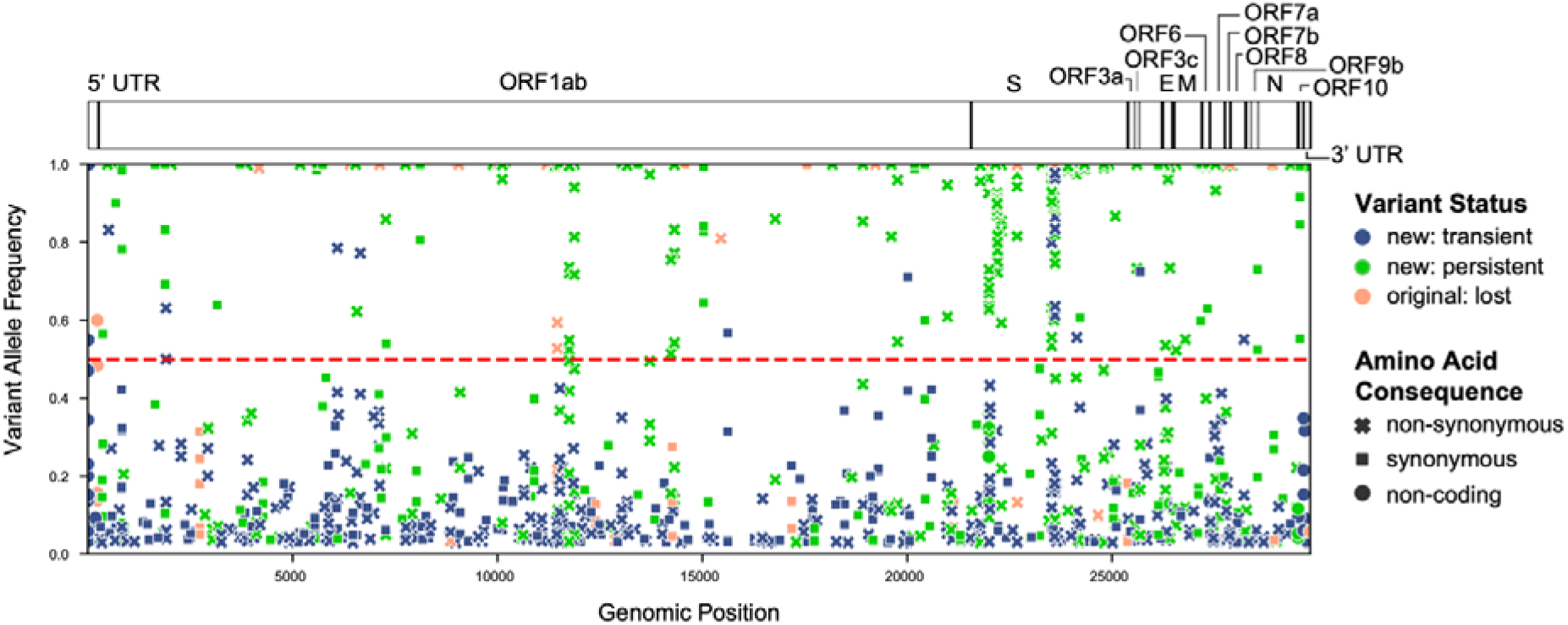
The location of all detected variants across all samples in all passage lines. The colour of points indicates whether a variant was present in the original clinical isolate (‘original: lost’), or arose new during serial passaging but was then lost (‘new: transient’) or persisted in the final passage number (‘new: persistent’). The shape of points corresponds to the inferred amino acid consequence, either non-synonymous, synonymous, or ‘other’ (in non-coding regions). All points above the horizontal dashed line are the majority allele for a given sample (allele frequency ≥50%). The location of genes within the SARS-CoV-2 genome is indicated above the plot in a schematic of the complete SARS CoV-2 genome of ∼30 kb.

**Figure 2:**
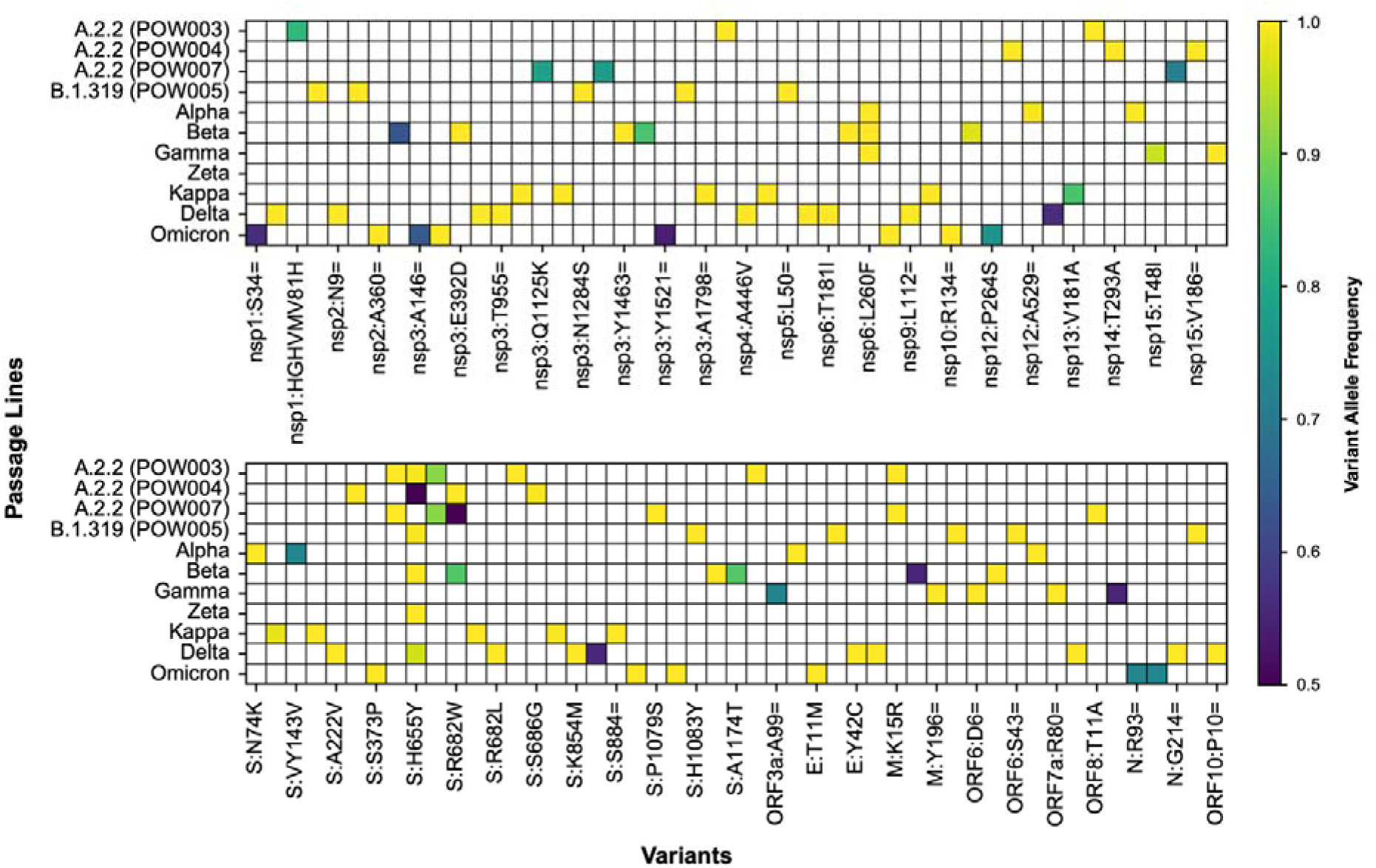
All variants that reached a consensus-level (>0.5) variant allele frequency (VAF) among passage lines, with coloured shading representing the maximum VAF reached within each given passage line. The notation for variant represents the inferred amino acid consequences of variants in different genes (see Supplementary Table S2 for further detail), with synonymous variants represented using ‘=’, as per HGVS Nomenclature recommendations.

**Figure 3:**
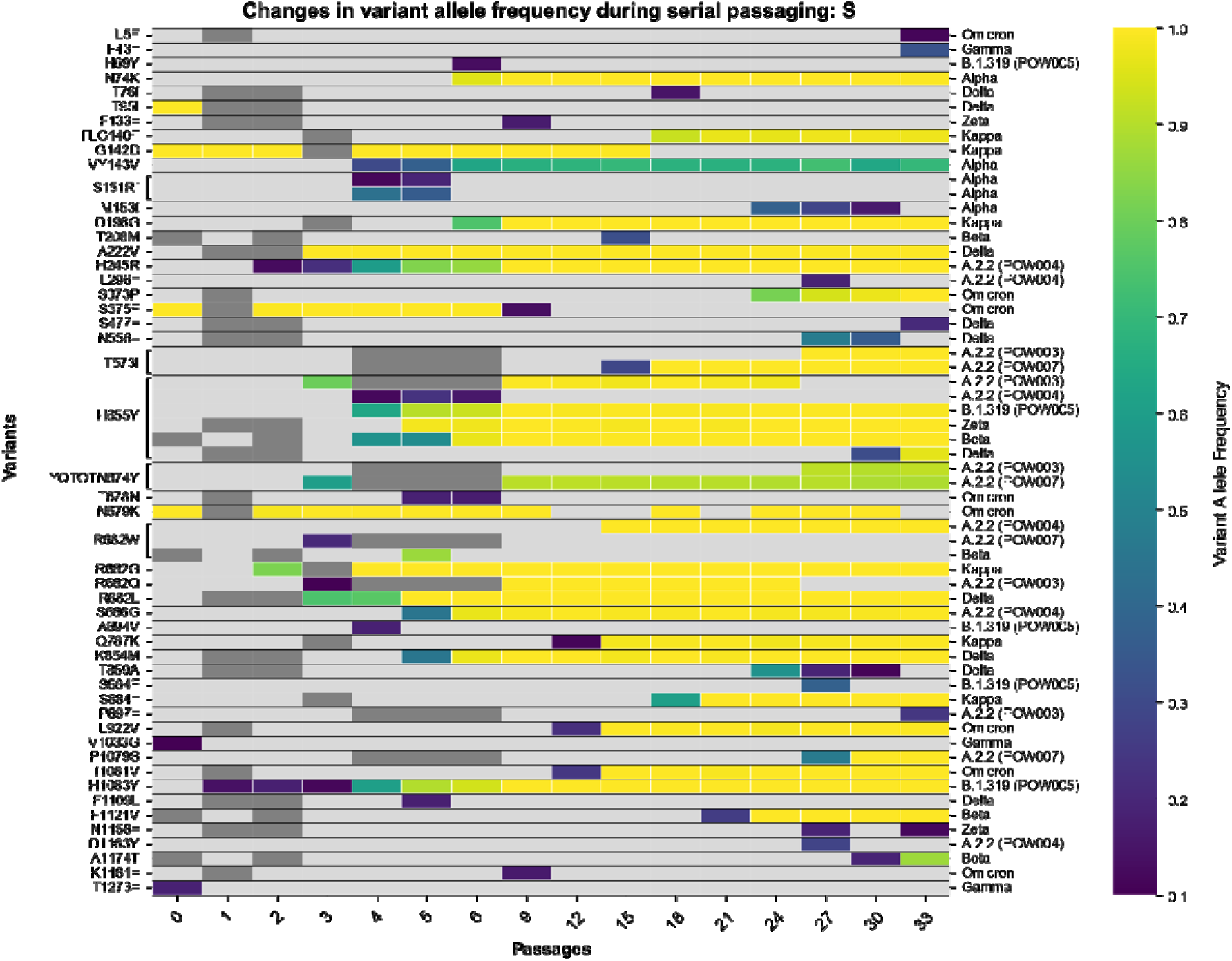
Heatmap depicting the variants detected in the spike gene of SARS-CoV-2 in each passage line throughout the course of serial passaging. The x-axis refers to the passage number (range: 0 [clinical isolate] to 33). On the primary y-axis the inferred amino acid consequences of variants in each gene are listed (see Supplementary Table S2 for further detail), with synonymous variants represented using ‘=’, as per HGVS Nomenclature recommendations. On the secondary y-axis (righthand side) the data in a given row is linked to the passage line from which it derives. Each ‘cell’ within the heatmap is shaded based on the variant allele frequency of a given variant in a given passage number within that passage line (light grey: undetected). Any cells that are dark grey represent cases where there is no sequencing data for that passage number. Convergence among passage lines is indicated by brackets linking a given variant to multiple passage lines. The dagger symbol represents a case where two different mutations at the nucleotide level led to the same inferred amino acid consequence.

**Figure 4:**
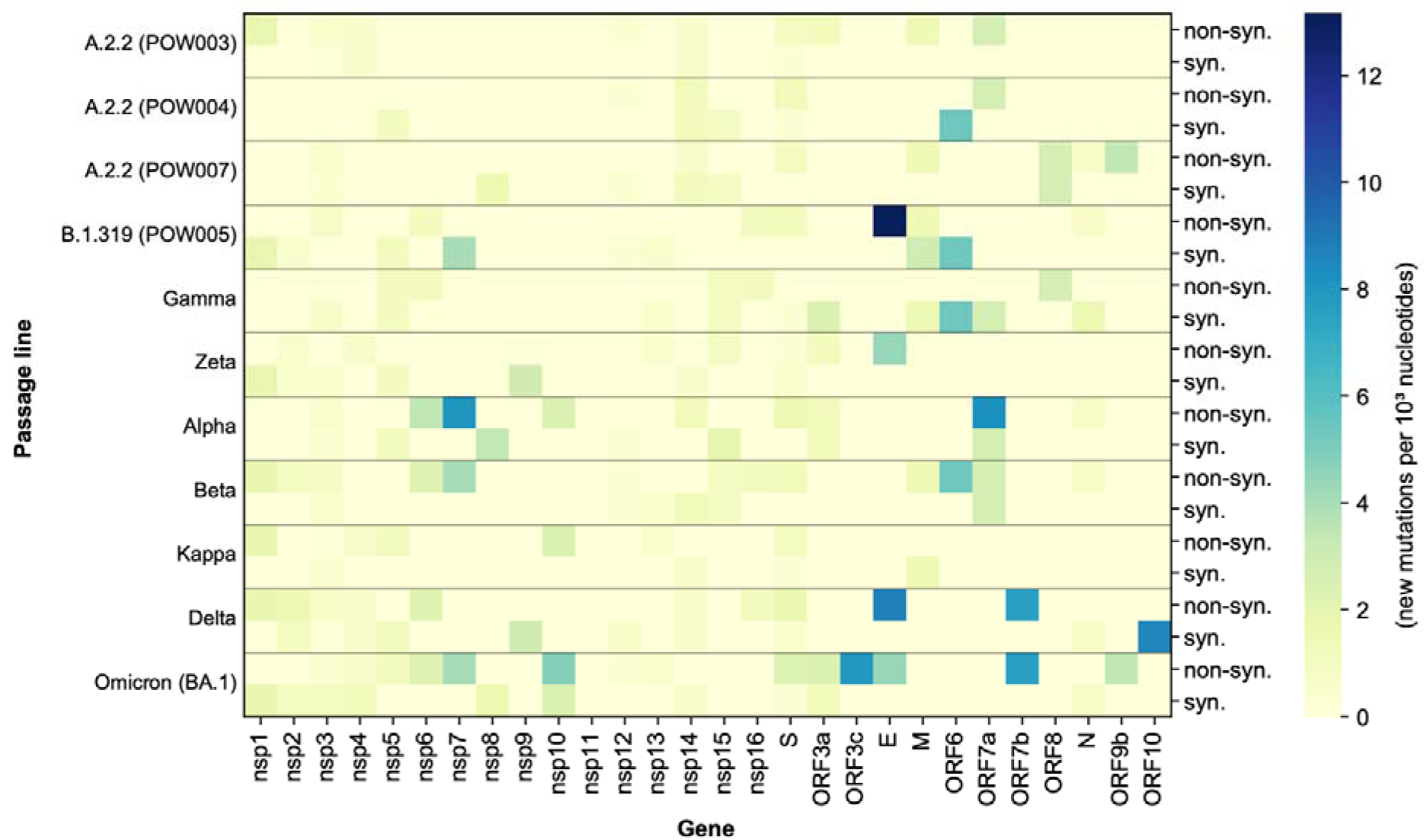
Heatmap depicting the total number of new mutations within each gene of the SARS-CoV-2 genome (normalised by gene length) throughout a maximum of 33 serial passages.

The accumulation of mutations over time was reflected in a gradual divergence between passage 0 and the final passage *within* each passage line, but without significantly affecting the phylogenetic relationships between passage lines (Figure 5). Nearly all samples within a given passage line form clades exclusive of the other passage lines, with the only exceptions being some poorly supported relationships among POW003 and POW007 (both of the A.2.2 Pango lineage). These results demonstrate that the convergent acquisition of some mutations among passage lines (as discussed below) did not impact upon inferred phylogenetic relationships. Overall, the phylogenetic relationships among passage lines reflects the expected topology based on the relationships among SARS-CoV-2 lineages within the global phylogeny. The inferred mutation rates *in vitro* varied among passage lines and varied within passage lines over time (Figure 6). There was a more than threefold difference in mutation rate between the passage lines with the greatest (Delta; ∼8.51 x 10^-05^ mutations per passage per nucleotide) and smallest (“POW004”; ∼2.64 x 10^-05^ mutations per passage per nucleotide) mutation rates. In general, across passage lines there was a trend to a reduced mutation rate as the number of serial passages increased.

**Figure 5:**
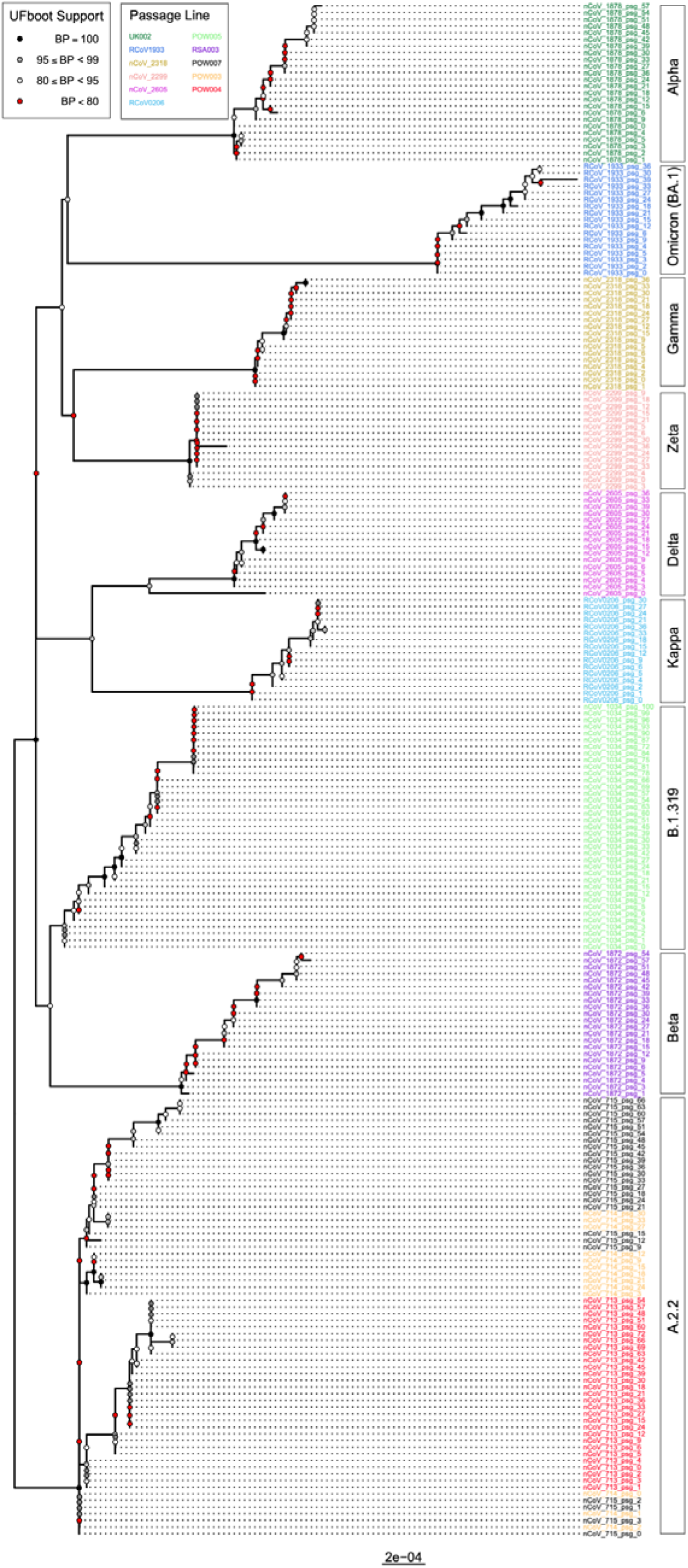
Phylogenetic tree depicting the inferred phylogenetic relationships among all samples in the study. The support for relationships (ultrafast bootstrap support, “UFboot”) is represented by shaded circles at nodes. Tip labels are shaded to reflect passage lines. The lineage calls for samples are depicted in rectangles to the right of the tree.

**Figure 6:**
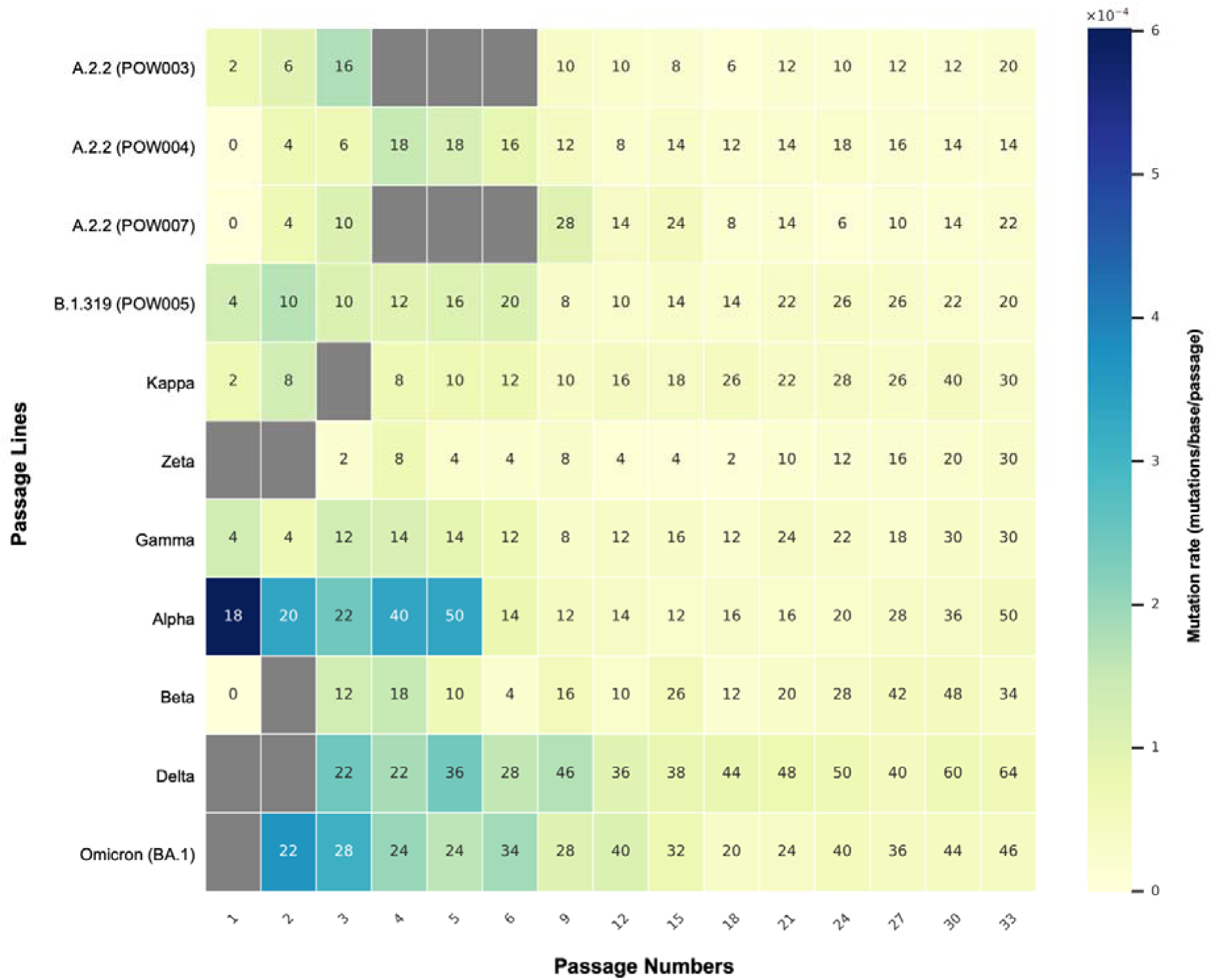
Heatmap depicting the inferred mutation rate (mutations per base per passage) at each passage number within each passage line. The rate was calculated based on the number of de novo mutations detected in a given passage number relative to the previous passage number (see main text). Here, “mutation” refers to any variant (SNP or indel) detected at a site or range of sites that pass all quality control metrics in a given sample. Each ‘cell’ in the heatmap is shaded based on the inferred mutation rate, and numbers within the cell represent the total number of detected mutations in that sample relative to the Wuhan Hu-1 reference genome for SARS-CoV-2. The purging of mutations (often those at a low variant allele frequency) during serial passaging can be observed, for example when comparing passages five and six in the Alpha passage line.

### No plateau after 100 serial passages

The “POW005” (Pango lineage: B.1.319) passage line was carried through the greatest number of passages across all passage lines in this study. There was a continuous accumulation of mutations in this passage line throughout the course of serial passaging, even up to passage 100 (Figure 7, Supplementary Table S2). The initial (passage 0) isolate possessed 10 variants relative to the Wuhan Hu-1 reference genome: one in the 5’ UTR and nine occurring in coding regions, of which eight were present as the majority allele and seven were non-synonymous variants. Throughout the course of serial passaging, POW005 accumulated 131 new variants (58 missense including two in-frame deletions, 63 synonymous, 10 in intergenic regions/UTRs), of which 97 were transient and 34 persisted until passage 100. The sole low-frequency variant detected in passage 0 was lost during passaging, although the other 10 variants present at passage 0 were retained. The passage line was still readily replicating with no obvious signs of impaired fitness.

**Figure 7:**
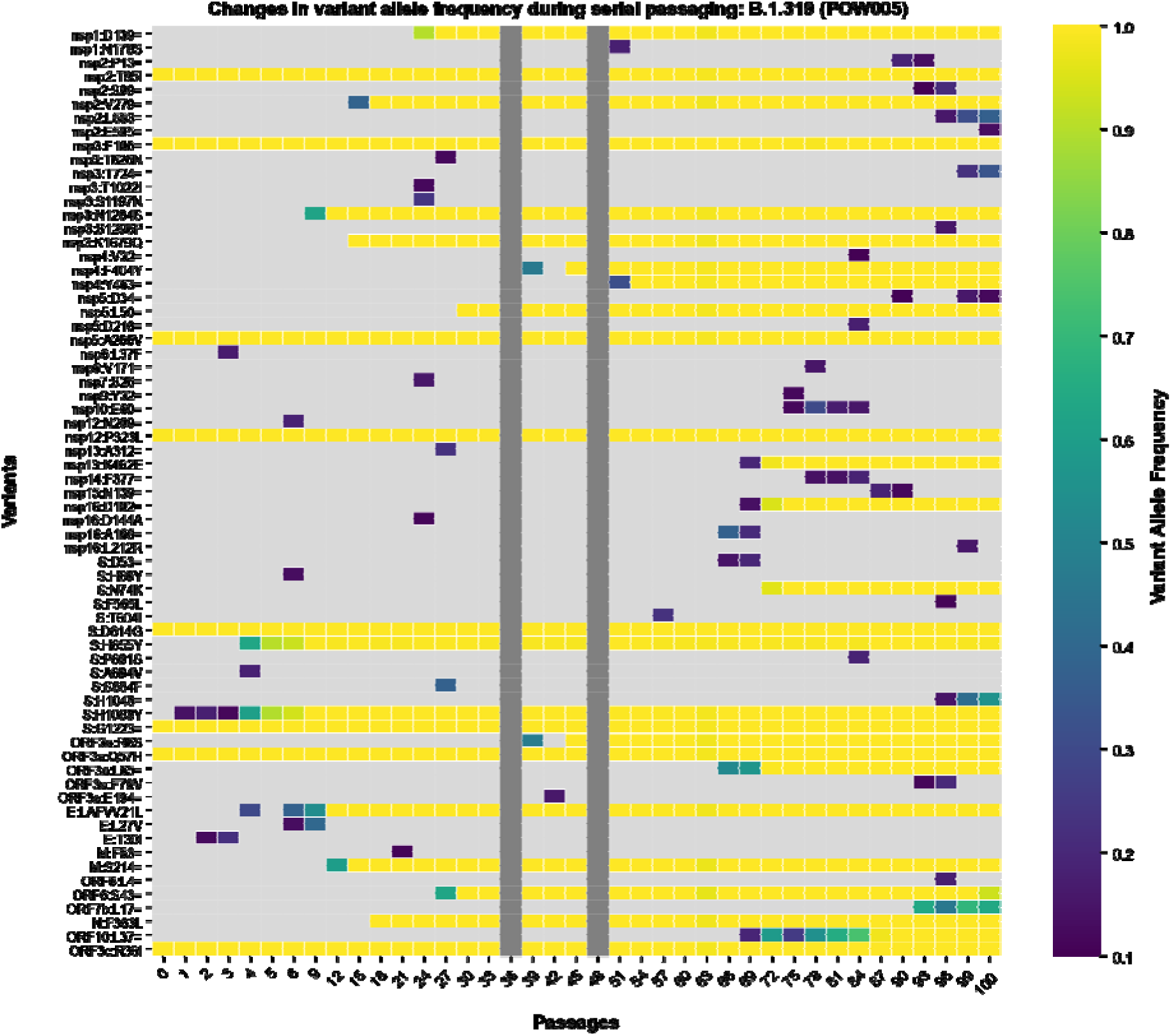
Heatmap depicting the variants detected in the each gene of SARS-CoV-2 in the POW005 passage line throughout the course of 100 serial passages. The x-axis refers to the passage number (range: 0 [clinical isolate] to 100). On the y-axis the inferred amino acid consequences of variants in each gene are listed (see Supplementary Table S2 for further detail), with synonymous variants represented using ‘=’, as per HGVS Nomenclature recommendations. Each ‘cell’ within the heatmap is shaded based on the variant allele frequency of a given variant in a given passage number within that passage line (light grey: undetected). Any cells that are dark grey represent cases where there is no sequencing data for that passage number.

### Mutations by gene location

Previous studies have concluded that the spike region of SARS-CoV-2 is a mutational hotspot. Whilst we observed many mutations arising in the spike gene (Figure 3), after correcting for gene length, the number of mutations in the spike gene was comparable with that of other genes (Figure 4). Genes coding for other structural and accessory proteins (e.g. M, ORF7a, ORF7b) accumulated mutations at a greater rate than other regions, including spike, with variation between passage lines (Figure 4). While the spike region is clearly an important region of the SARS-CoV-2 genome for mutation (Figure 3), our results further reflect recent studies highlighting the importance of other regions of the genome for understanding the ongoing evolution of SARS-CoV-2 (42–45), and the capacity of the virus to adapt to new evolutionary pressures.

### Signatures of convergent evolution

As the COVID-19 pandemic progressed, multiple mutations speculated to impart fitness benefits on SARS-CoV-2 were observed to have arisen multiple times independently. Examples include the spike mutations N501Y, E484K and ΔH69/V70, which increased immune escape potential and/or increased ACE2 binding affinity (7, 9, 46–48). Therefore, to understand further the impact of the mutations arising *de novo* during long-term passaging, we determined whether there were any signatures of convergent evolution. Although most variants arising during serial passaging were passage-line specific, there were also many cases where different passage lines independently acquired the same variants convergently (Figure 8). In total, there were 12 variants that arose throughout serial passaging whereby (a) the variant was present in two or more passage lines, and (b) it newly arose in at least one of the passage lines (Table 2). Occasionally these went to fixation in multiple passage lines. Convergent mutations are known to arise during serial passaging in Vero E6 cells, especially in the region of the spike furin-cleavage site (49, 50). We found that the S:YQTQTN674Y mutation, a deletion of five amino acids near the spike S1/S2 cleavage site, arose independently in POW003 (A.2.2) and POW007 (A.2.2). This QTQTN motif has been experimentally linked to pathogenesis, with its loss in clinical and passaged viruses leading to attenuated viral replication and disease, driven by less efficient spike processing through hindered cleavage of the furin-cleavage site (15, 51–54). We also observed four different missense amino acid replacements proximal to the furin-cleavage site at amino acid position 682 of spike (R682G, R682W, R682L, R682Q) at varying allele frequencies, as well as a six amino acid deletion in one passage line (RARSVA683R). All but one of these mutations appeared within four serial passages.

**Figure 8:**
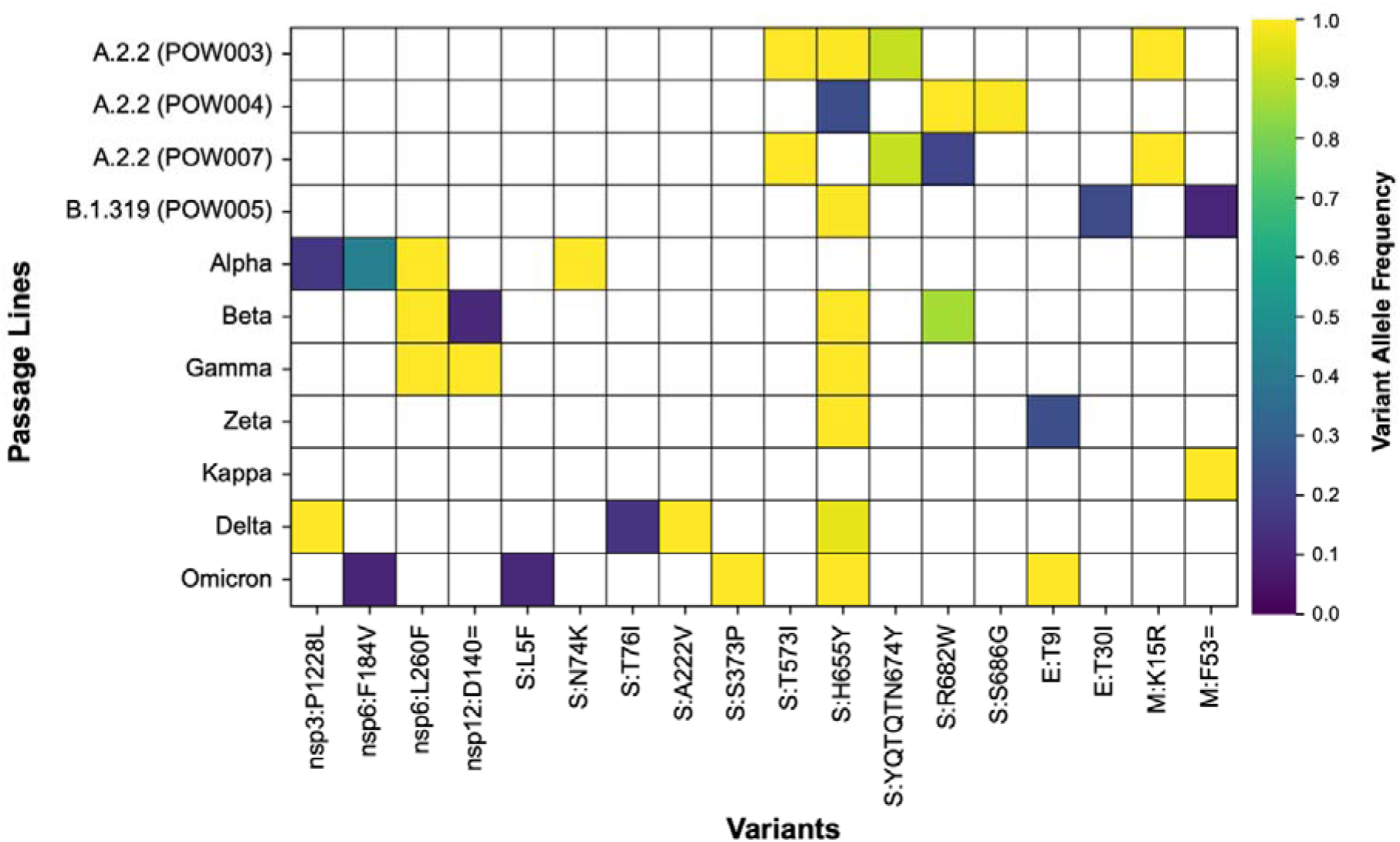
All variants that arose *de novo* during serial passaging in at least one passage line and were (a) also detected in at least one other passage line, and/or (b) present in a curated compendium of functional impacts of SARS-CoV-2 variants (see main text). These variants represent cases of convergent evolution. Coloured shading depicts the maximum variant allele frequency reached by each variant in each passage line.

**Table 2:**
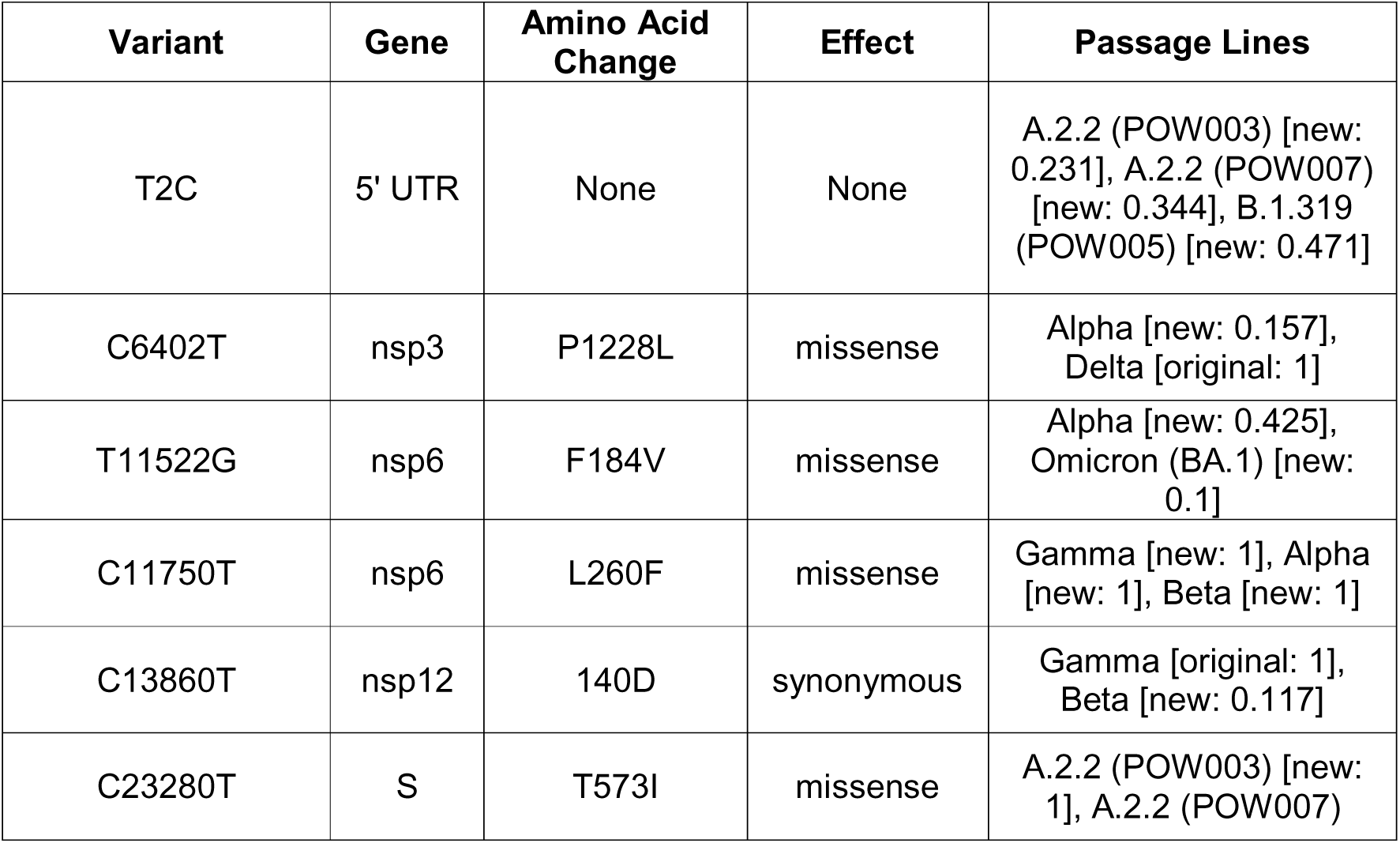
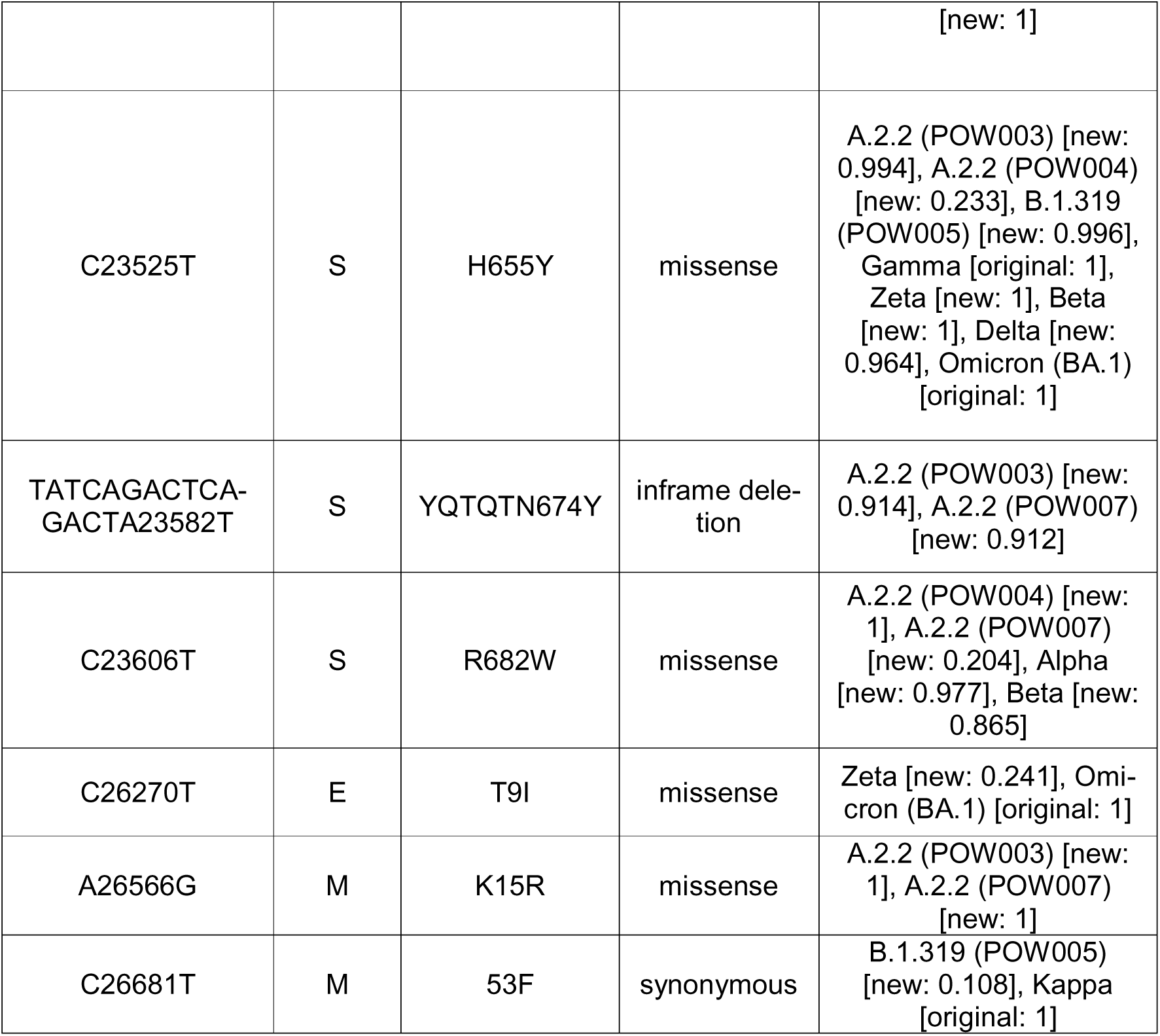
All variants that arose *de novo* during the course of *in vitro* serial passaging that were also present (either originally or newly arising) in at least one other passage lines. The numbers in square brackets in the ‘lines’ column refer to the maximum variant allele frequency that each variant was detected at in each corresponding passage line.

Any mutations associated with enhanced virus replication *in vitro* would be expected to be beneficial for virus survival and selected for, as is the case with replication-enhancing mutations in the clinical population. For example, the emergence of the spike mutation D614G early in the COVID-19 pandemic was linked with the proliferation of virus with enhanced replication and transmission (55–57), as well as changes in confirmation of the spike protein (6, 58). Additional changes in the spike region have been linked to enhanced virus replication at subsequent stages of the pandemic, such as the P681R mutation within the Delta VOC (59, 60). However, the impact of mutations in other genomic regions on virus replication, including replication fidelity, is also important to consider. We focused initially on *de novo* mutations in nsp5 (protease, 3CL^pro^), nsp10 (cofactor that stimulates ExoN), nsp12 (RNA-dependent RNA polymerase, RdRp), and nsp14 (exonuclease, ExoN) given the important roles these non-structural proteins play in replication fidelity. There were 15 non-synonymous mutations in these regions, some of which became the majority allele and/or went to fixation (Table 3). The nsp14:P203L mutation, which arose at a low VAF in passage line POW007, has been (a) suggested to lead to an increased evolutionary rate *in vivo* (61), and (b) linked to altered replication over 250 passages of murine hepatitis virus (62). Other mutations in Table 3 have not been explicitly linked to altered replication, but have been temporally associated with resistance to antivirals that target these replication-associated non-structural proteins (63, 64).

**Table 3:**
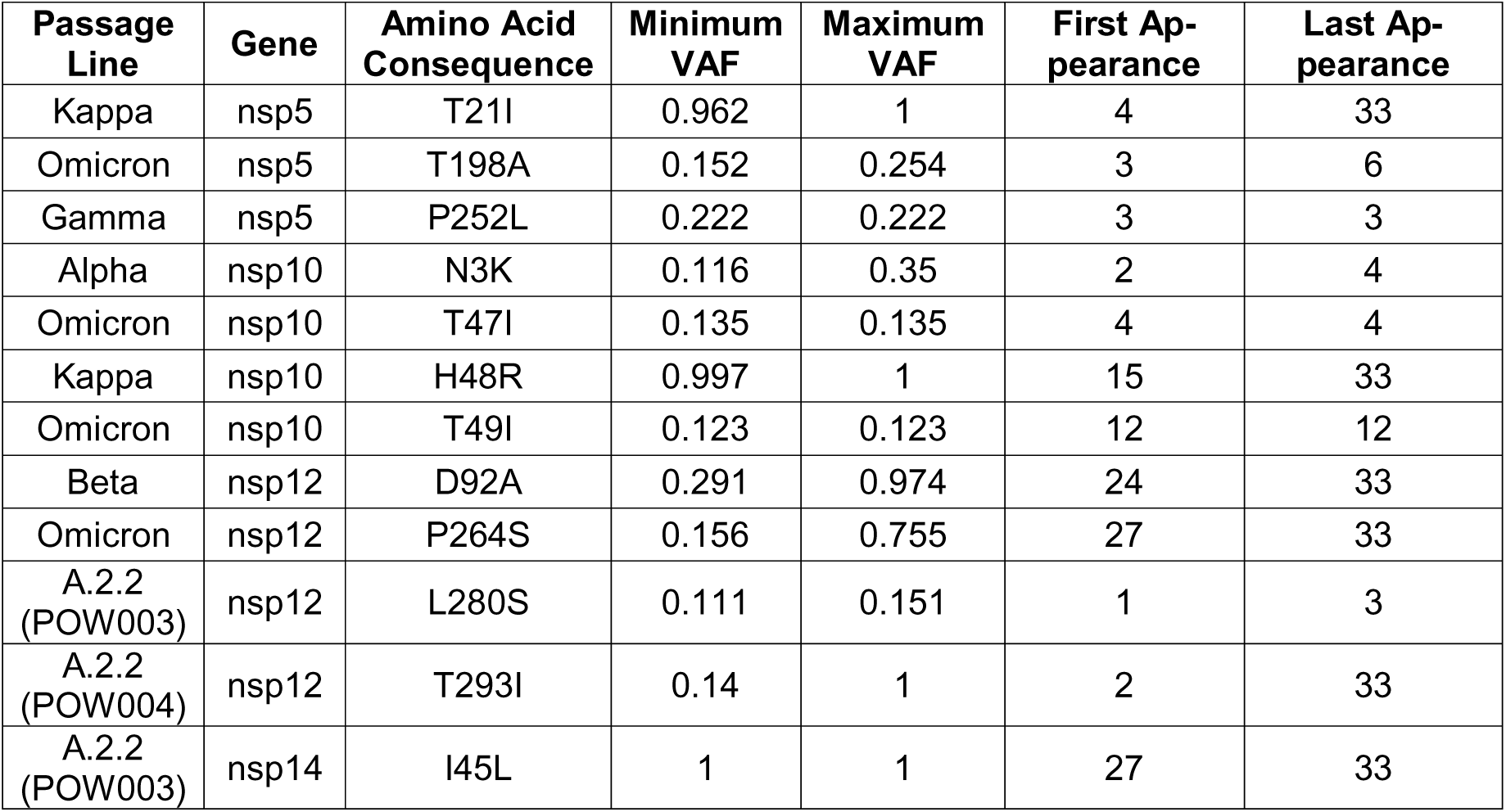
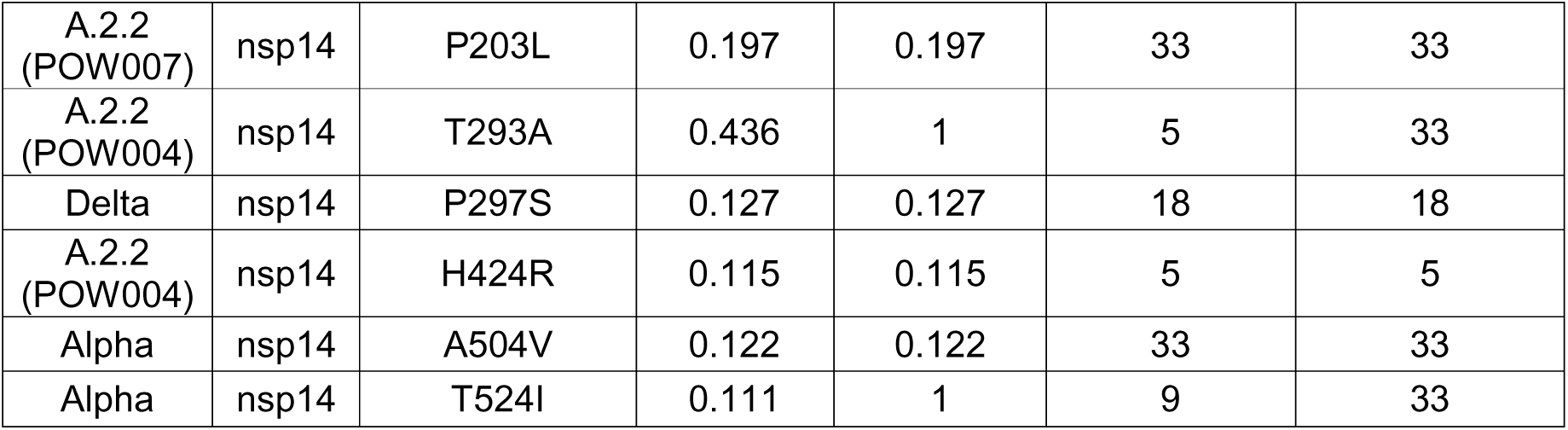
All mutations arising *de novo* throughout the course of long-term serial passaging that fall within nsp5, nsp10, nsp12, and nsp14 of the SARS-CoV-2 genome.

| Passage Line | Gene | Amino Acid Consequence | Minimum VAF | Maximum VAF | First Appearance | Last Appearance |
| --- | --- | --- | --- | --- | --- | --- |
| Kappa | nsp5 | T21I | 0.962 | 1 | 4 | 33 |
| Omicron | nsp5 | T198A | 0.152 | 0.254 | 3 | 6 |
| Gamma | nsp5 | P252L | 0.222 | 0.222 | 3 | 3 |
| Alpha | nsp10 | N3K | 0.116 | 0.35 | 2 | 4 |
| Omicron | nsp10 | T47I | 0.135 | 0.135 | 4 | 4 |
| Kappa | nsp10 | H48R | 0.997 | 1 | 15 | 33 |
| Omicron | nsp10 | T49I | 0.123 | 0.123 | 12 | 12 |
| Beta | nsp12 | D92A | 0.291 | 0.974 | 24 | 33 |
| Omicron | nsp12 | P264S | 0.156 | 0.755 | 27 | 33 |
| A.2.2 (POW003) | nsp12 | L280S | 0.111 | 0.151 | 1 | 3 |
| A.2.2 (POW004) | nsp12 | T293I | 0.14 | 1 | 2 | 33 |
| A.2.2 (POW003) | nsp14 | I45L | 1 | 1 | 27 | 33 |
| A.2.2 (POW007) | nsp14 | P203L | 0.197 | 0.197 | 33 | 33 |
| A.2.2 (POW004) | nsp14 | T293A | 0.436 | 1 | 5 | 33 |
| Delta | nsp14 | P297S | 0.127 | 0.127 | 18 | 18 |
| A.2.2 (POW004) | nsp14 | H424R | 0.115 | 0.115 | 5 | 5 |
| Alpha | nsp14 | A504V | 0.122 | 0.122 | 33 | 33 |
| Alpha | nsp14 | T524I | 0.111 | 1 | 9 | 33 |

A further literature search revealed that at least 16 different variants that arose *de novo* during serial passaging have important functional consequences in the global clinical population of SARS-CoV-2, as recorded under 54 functional categories (19 unique) in a curated compendium of scientific literature (35) (Table 3). Noted functional impacts of the newly arising mutations in our study, including the spike mutations L5F, A67V, T76I, A222V and others, are related to various aspects of pharmaceutical effectiveness, including vaccine efficacy and antibody neutralization (Table 4). The repeated occurrence of mutations *in vitro* through serial passaging that are congruent with those related to immune evasion, even in the absence of immune or drug-related selective pressures, suggests that at least some mutations in the clinical population might not have been driven by intra-host selection (cf. Weigang et al. (65)). That is, these clinical population mutations may arise *de novo* with intrinsic immune escape benefits, or there is another benefit of these putative immune-related mutations to the survival of SARS-CoV-2 separate to immune escape.

**Table 4:**
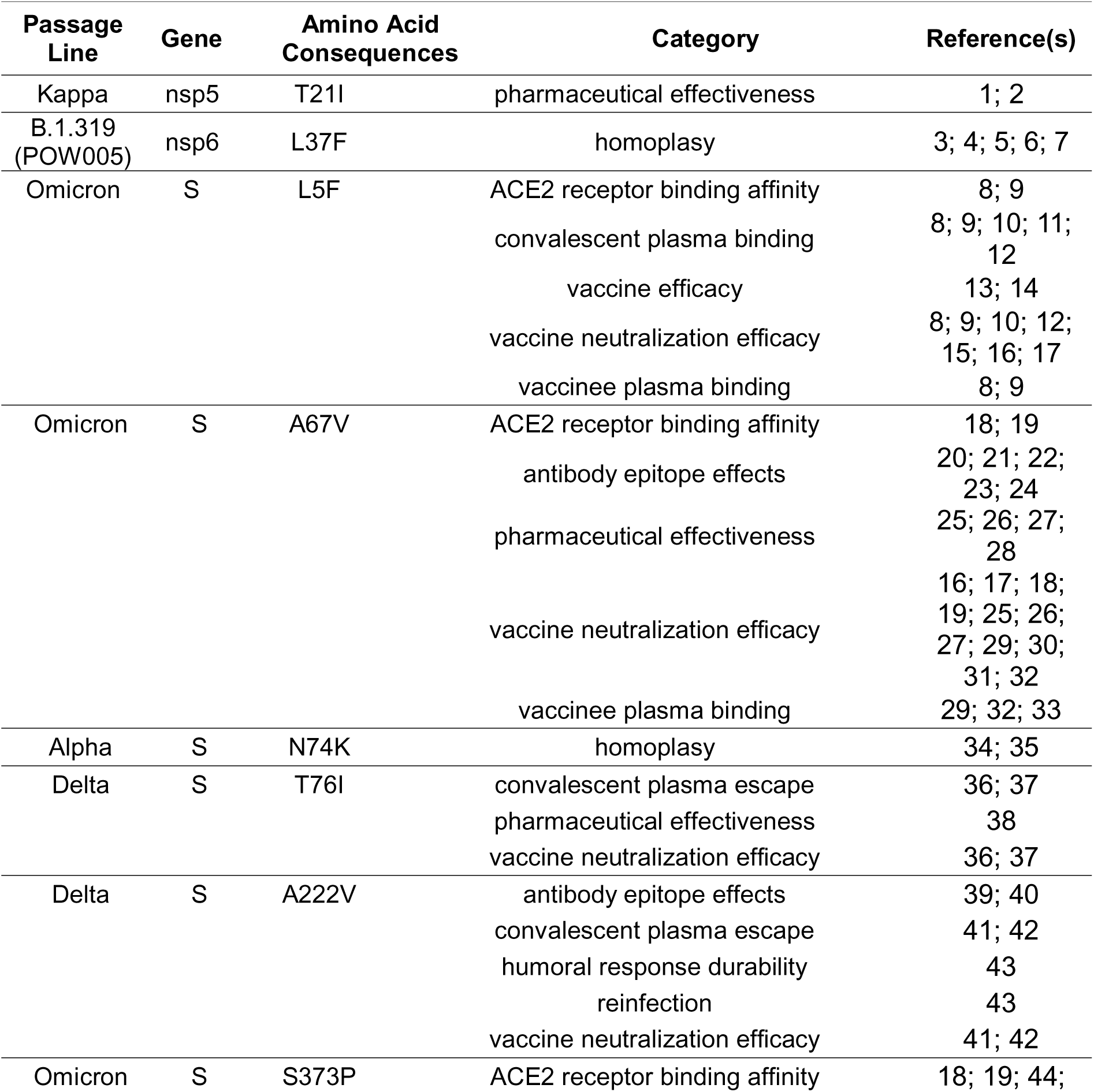
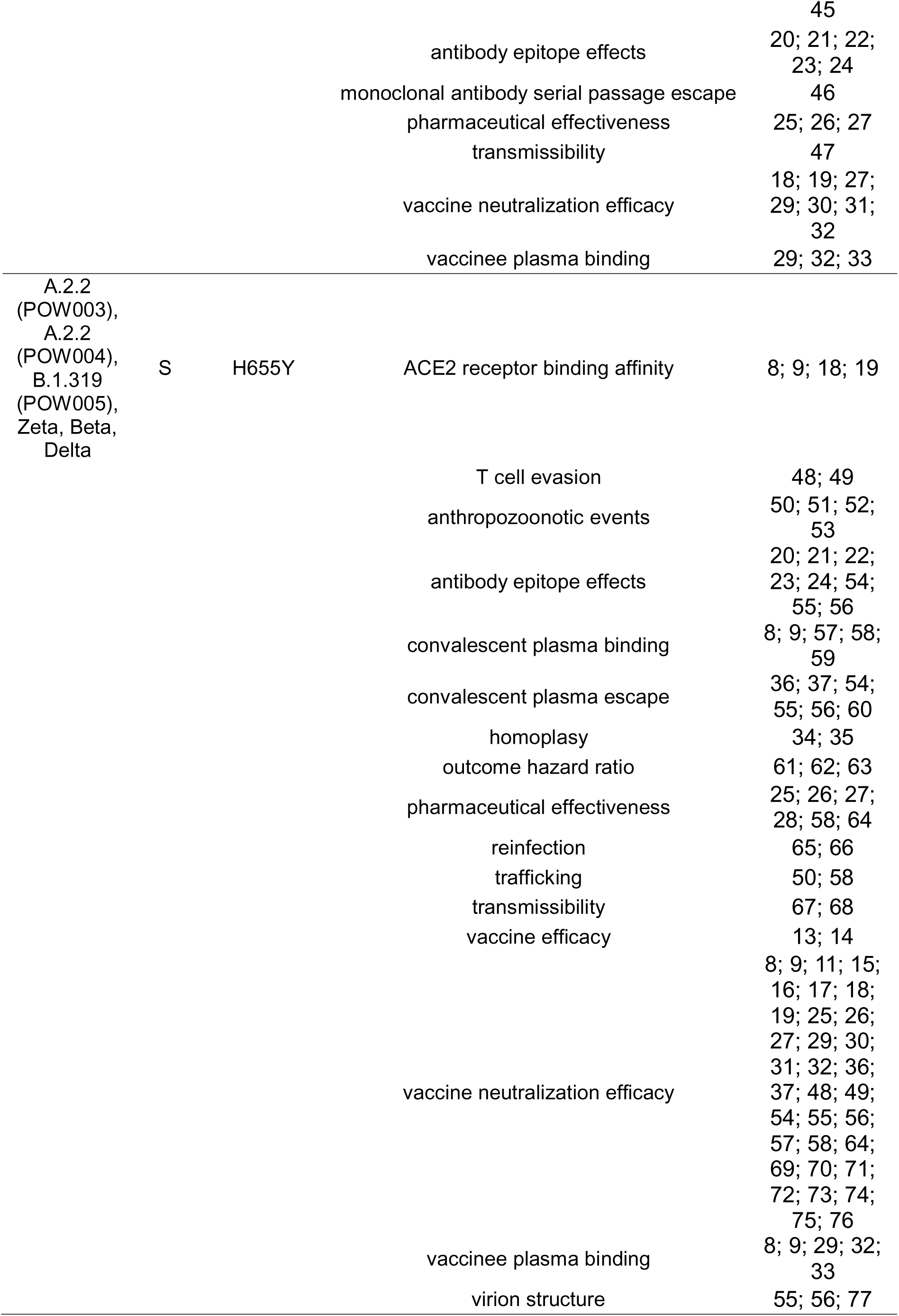
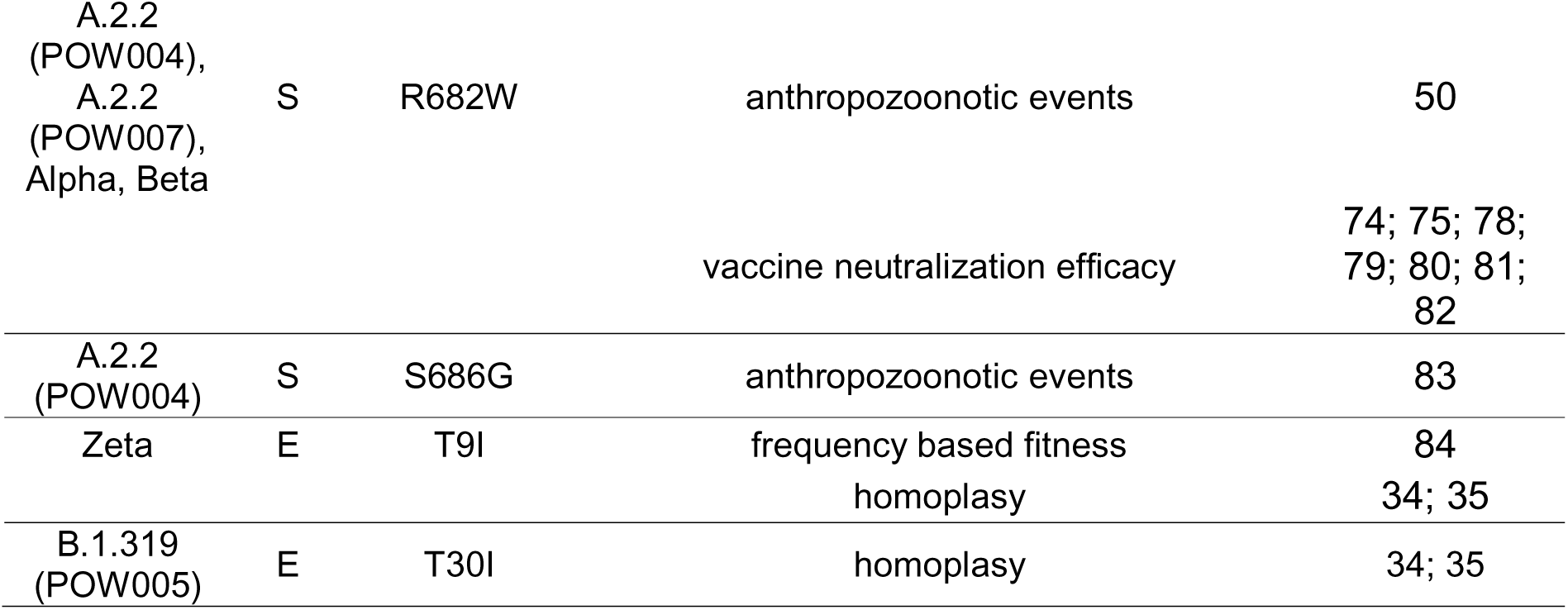
All mutations arising *de novo* throughout the course of long-term serial passaging that are noted in a compendium of functional impacts of mutations in the SARS-CoV-2 Wuhan Hu-1 reference genome (https://github.com/nodrogluap/pokay). All passage lines we studied (see text) are noted in the first column. Reference numbers refer to those in Supplementary File S1.

We also observed the loss of at least one clinically significant mutation throughout the course of serial passaging. One key variant hypothesised to have contributed to the success of the Omicron BA.1 lineage is S:S375F, a mutation within the receptor-binding domain of the Spike protein responsible for attenuation of spike cleavage efficiency and fusogenicity as well as decreased ACE2 binding affinity (66). This key mutation was present in our BA.1 passage line at a VAF of ∼1 from passage 0 until passage 6 (i.e., fixed as the majority allele), but was only present at a low VAF of 13% in passage 9, and was then not detected in subsequent passages (Supplementary Table S2). The dominance of S:S375F in global Omicron sequences suggests the site may have been subject to initial positive selection followed by purifying selection to retain benefits imparted by the mutation, or may represent a form of bottleneck.

### A putative rapid switch in cell entry pathway *in vitro*

Aside from the deletion of the QTQN motif (see above), the most striking example of convergence across passage lines was the repeated *de novo* evolution of S:H655Y (Figures 2–3, 8). This mutation was already present in passage 0 in two passage lines and arose newly throughout serial passaging in six of the remaining nine passage lines. The S:H655Y mutation was originally detected in early cases of the Gamma and Omicron VOCs. Impacts of the S:H655Y mutation include acting as a “gateway” mutation by increasing the fitness of lineages with complex combinations of clinically significant mutations, and increased spike cleavage and fusogenicity associated with enhancement of the endosomal cell entry pathway (6, 67, 68). A putative switch towards preferentially using an endosomal cell entry pathway within the Vero E6 passage lines, rather than the ACE2-TMPRSS2 pathway typical of most variants prior to Omicron (69), is not surprising given that Vero E6 cell lines do not express TMPRSS2. Similar changes in virus tropism have been observed in the clinical population. For example, several lineages of the Omicron BA.1 and BA.2 variants (and their descendants) used an endosomal cell entry pathway rather than the ACE2-TMPRSS2 pathway for virus entry (70). This change in tropism resulted in Omicron replicating more efficiently in the bronchi, but not productively infecting human alveolar type 2 cells of the lung (71, 72).

### Adaptation, bottlenecks and genetic drift

We observed many variants newly arising in all passaged SARS-CoV-2 lineages (Figures 1–3, 7– 9; Supplementary Figures S1–26; Supplementary Table S2). Some of the >170 newly arising variants were transient and purged from the virus populations throughout the course of serial passaging, but some persisted until the final serial passages, occasionally at (near-) fixation within the virus populations. One signature of positive selection is a selective sweep, whereby a rare or previously non-existing allele rapidly increases in frequency within a population. Therefore, we considered the fixation of these newly arising mutations throughout the course of serial passaging to represent possible positive selection for mutations beneficial to virus survival *in vitro.* Examples include the furin cleavage site-associated mutations and mutations associated with a change in cell entry pathway (discussed above).

**Figure 9:**
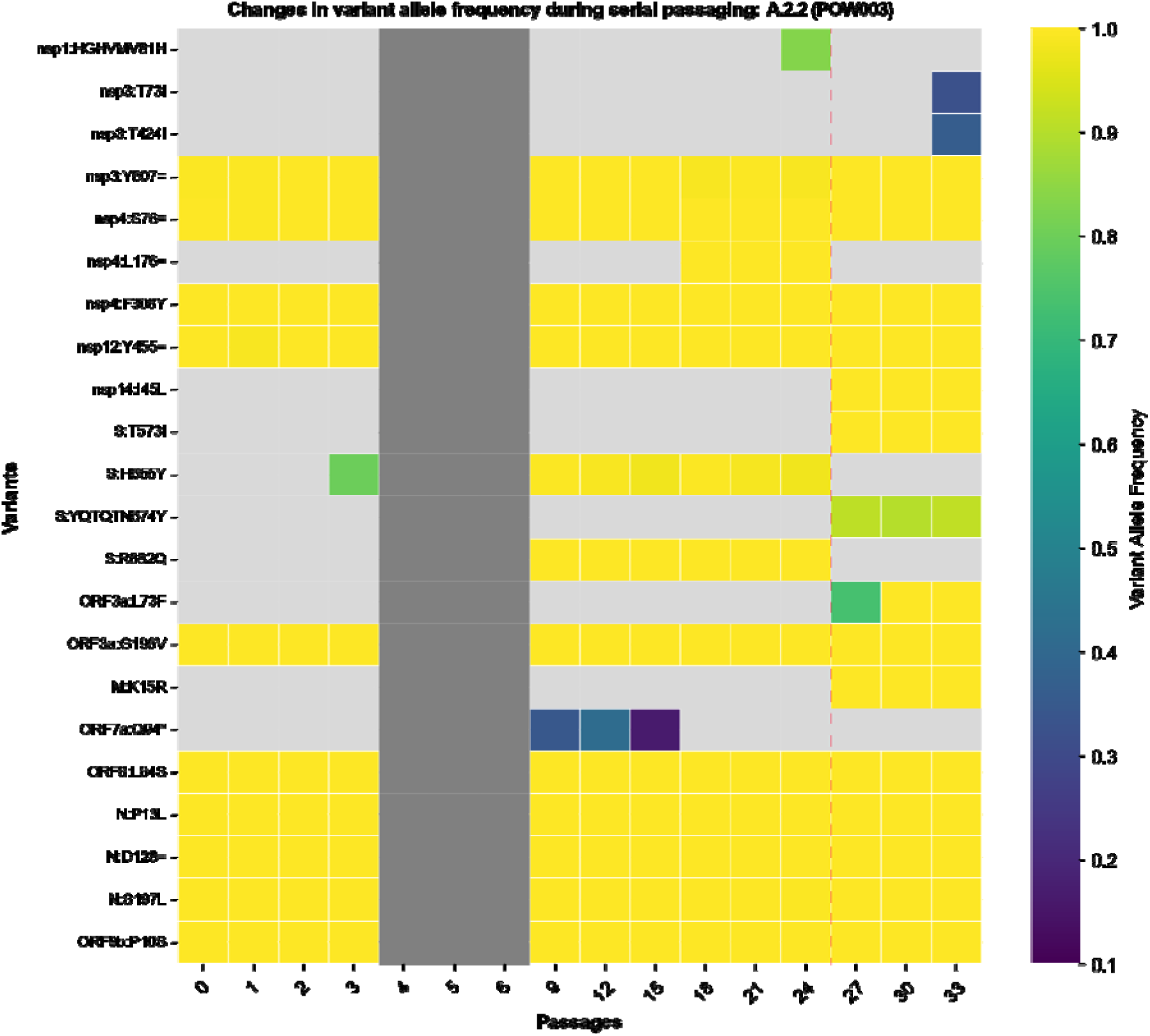
Heatmap depicting the variants detected in the each gene of SARS-CoV-2 in the POW003 passage line throughout the course of 33 serial passages. The x-axis refers to the passage number (range: 0 [clinical isolate] to 33). On the y-axis the inferred amino acid consequences of variants in each gene are listed (see Supplementary Table S2 for further detail), with synonymous variants represented using ‘=’, as per HGVS Nomenclature recommendations. Each ‘cell’ within the heatmap is shaded based on the variant allele frequency of a given variant in a given passage number within that passage line (light grey: undetected). Any cells that are dark grey represent cases where there is no sequencing data for that passage number. The vertical dashed line represents a boundary point at which a partial haplotype replacement is inferred to have occurred (between passages 24 and 27) given the abrupt appearance of several fixed/high-frequency variants.

Although many mutations we observed have previously been linked to increased transmissibility and replication *in vivo*, their proliferation *in vitro* could instead be through chance alone, such as through transmission bottlenecks between successive serial passages. Our serial passaging protocol was designed to maintain an estimated MOI of 0.01, allowing for cytopathic effects to become evident within a reasonable time frame before passaging continued. Maintaining a lower virus titre during serial passaging can increase the impact of genetic bottlenecks, influencing the fate of emerging variants and potentially obscuring whether changes in variant frequency were driven by adaptation or stochastic events like genetic drift (73–75). One putative case of an abrupt shift in variant allele frequencies caused by a genetic bottleneck was observed in the POW003 (A.2.2), where an abrupt loss and gain of several variants occurred between passages 24 and 27 (Figure 9). Analogous situations in the global clinical populations of SARS-CoV-2 have been observed, where mutations that were either non-beneficial, or possibly deleterious, became fixed within given regions after public health measures created transmission bottlenecks (e.g. (43)). Likewise, the ORF8 accessory region remains a hotspot for mutations throughout the COVID-19 pandemic (76, 77), possibly as a result of a relaxation of purifying selection, as was hypothesised for SARS-CoV (78).

Regardless of the precise evolutionary mechanisms driving mutations in our study, our results and data represent a useful resource to the virology community. Apart from revealing signatures of convergence with previously noted mutations of interest, our results provide a resource for investigating future changes occurring in SARS-CoV-2 globally. Any mutations arising convergently among passage lines that have not been observed commonly as occurring during serial passaging, or in the global clinical population, are worth further investigation. The repeated appearance of these mutations during in vitro culture implies they benefit SARS-CoV-2, at least in vitro, and could also add to the growing list of Vero E6-associated mutations in SARS-CoV-2. The same is true for other mutations arising *de novo* even in a single passage line.

## Data availability

The consensus genomes of all passage 0 samples from this study have been uploaded to GISAID (see Table 1), and all sequence data have been uploaded to NCBI’s GenBank Sequence Read Archive under accession number PRJNA1018257. The pipeline to process and analyse raw sequencing reads is available from https://github.com/charlesfoster/covid-illumina-snakemake, and the pipeline to summarise longitudinal variant calling results is available from https://github.com/charlesfoster/vartracker.

