## Supplementary material for "Long-term serial passaging of SARS-CoV-2 reveals signatures of convergent evolution": Supplementary_Figures.pdf

Fig. S1

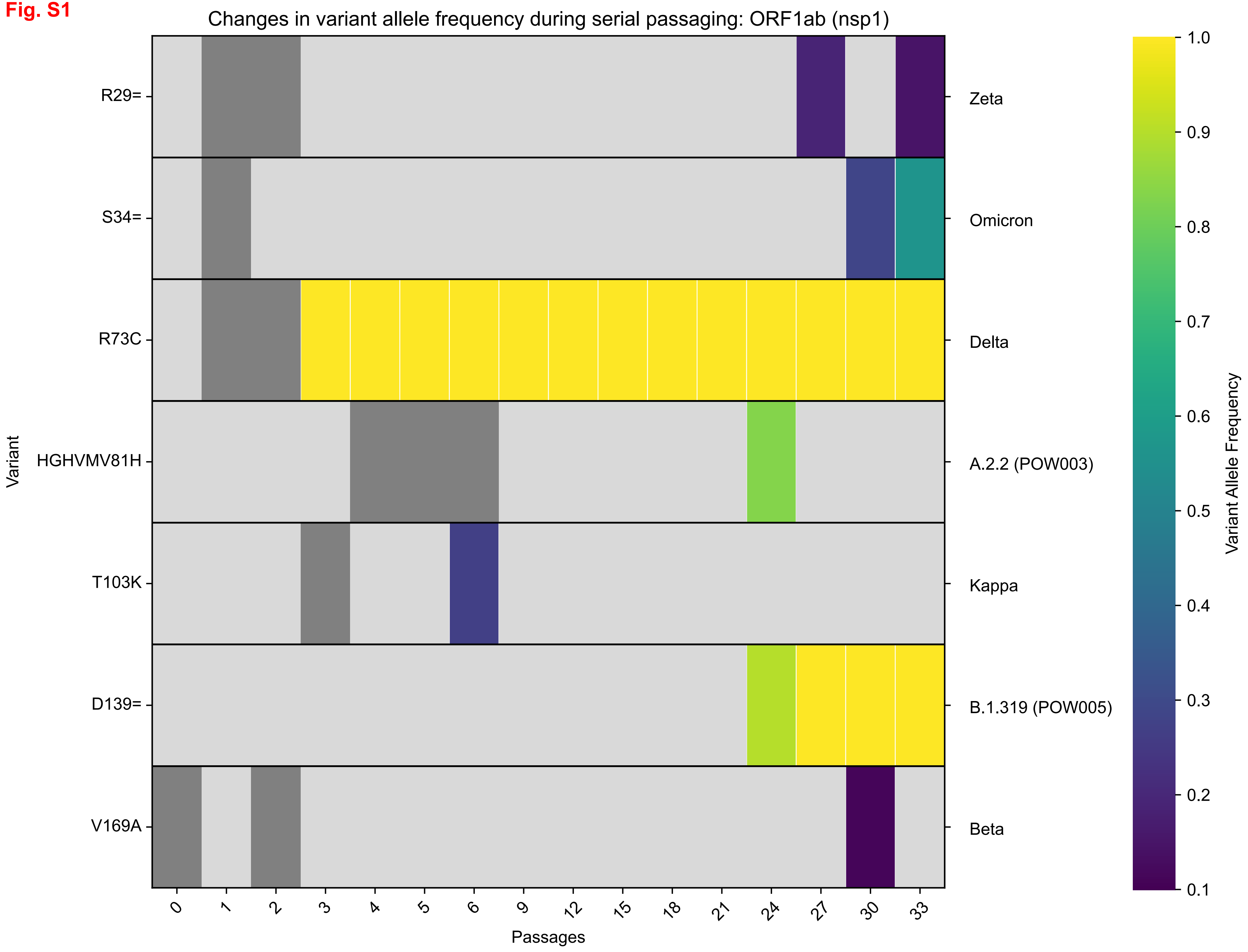

Changes in variant allele frequency during serial passaging: ORF1ab (nsp1)
Variant
R29=
S34=
R73C
HGHVMV81H
T103K
D139=
V169A
Zeta
Omicron
Delta
A.2.2 (POW003)
Kappa
B.1.319 (POW005)
Beta
0
1
2
3
4
5
6
9
12
15
18
21
24
27
30
33
Passages
Variant Allele Frequency
1.0
0.9
0.8
0.7
0.6
0.5
0.4
0.3
0.2
0.1

Fig. S2

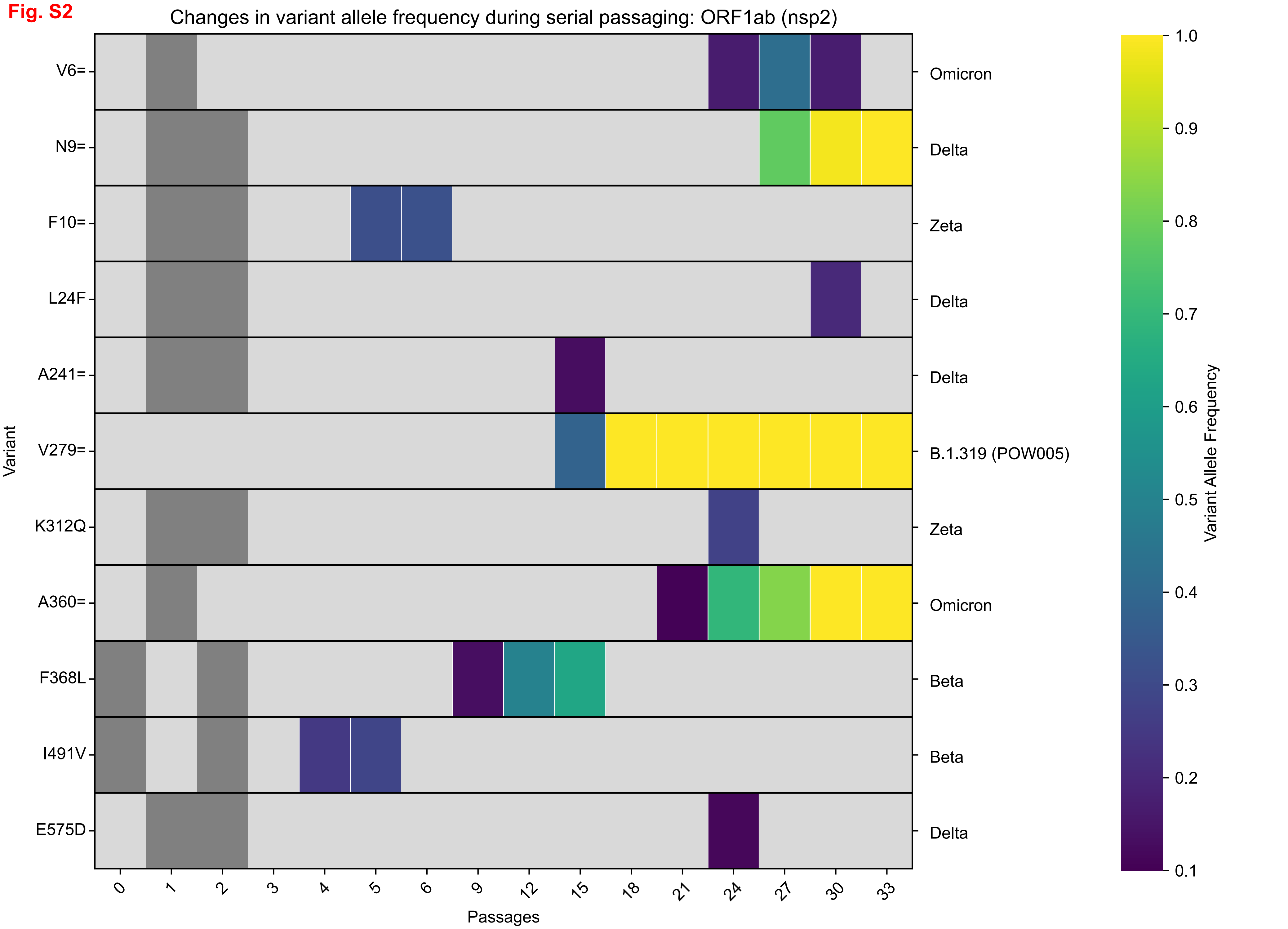
Changes in variant allele frequency during serial passaging: ORF1ab (nsp2)
Variant
V6=
N9=
F10=
L24F
A241=
V279=
K312Q
A360=
F368L
I491V
E575D
Omicron
Delta
Zeta
Delta
Delta
B.1.319 (POW005)
Zeta
Omicron
Beta
Beta
Delta
0
1
2
3
4
5
6
9
12
15
18
21
24
27
30
33
Passages
Variant Allele Frequency
1.0
0.9
0.8
0.7
0.6
0.5
0.4
0.3
0.2
0.1

Fig. S3

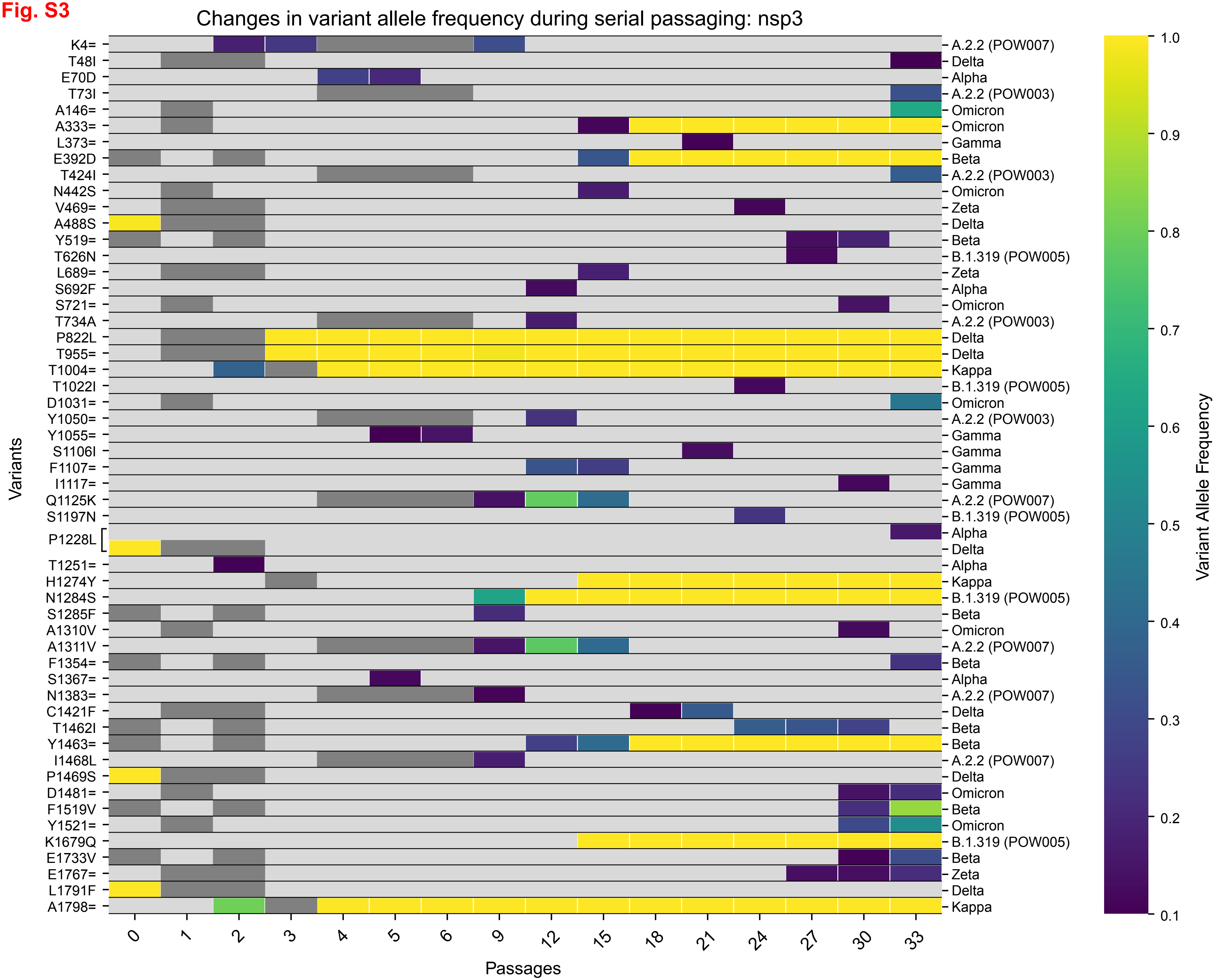

Changes in variant allele frequency during serial passaging: nsp3
Variants
K4=
T48I
E70D
T73I
A146=
A333=
L373=
E392D
T424I
N442S
V469=
A488S
Y519=
T626N
L689=
S692F
S721=
T734A
P822L
T955=
T1004=
T1022I
D1031=
Y1050=
Y1055=
S1106I
F1107=
I1117=
Q1125K
S1197N
P1228L
T1251=
H1274Y
N1284S
S1285F
A1310V
A1311V
F1354=
S1367=
N1383=
C1421F
T1462I
Y1463=
I1468L
P1469S
D1481=
F1519V
Y1521=
K1679Q
E1733V
E1767=
L1791F
A1798=
A.2.2 (POW007)
Delta
Alpha
A.2.2 (POW003)
Omicron
Omicron
Gamma
Beta
A.2.2 (POW003)
Omicron
Zeta
Delta
Beta
B.1.319 (POW005)
Zeta
Alpha
Omicron
A.2.2 (POW003)
Delta
Delta
Kappa
B.1.319 (POW005)
Omicron
A.2.2 (POW003)
Gamma
Gamma
Gamma
Gamma
A.2.2 (POW007)
B.1.319 (POW005)
Alpha
Delta
Alpha
Kappa
B.1.319 (POW005)
Beta
Omicron
A.2.2 (POW007)
Beta
Alpha
A.2.2 (POW007)
Delta
Beta
Beta
A.2.2 (POW007)
Delta
Omicron
Beta
Omicron
B.1.319 (POW005)
Beta
Zeta
Delta
Kappa
0
1
2
3
4
5
6
9
12
15
18
21
24
27
30
33
Passages
Variant Allele Frequency
1.0
0.9
0.8
0.7
0.6
0.5
0.4
0.3
0.2
0.1

**Fig. S4**

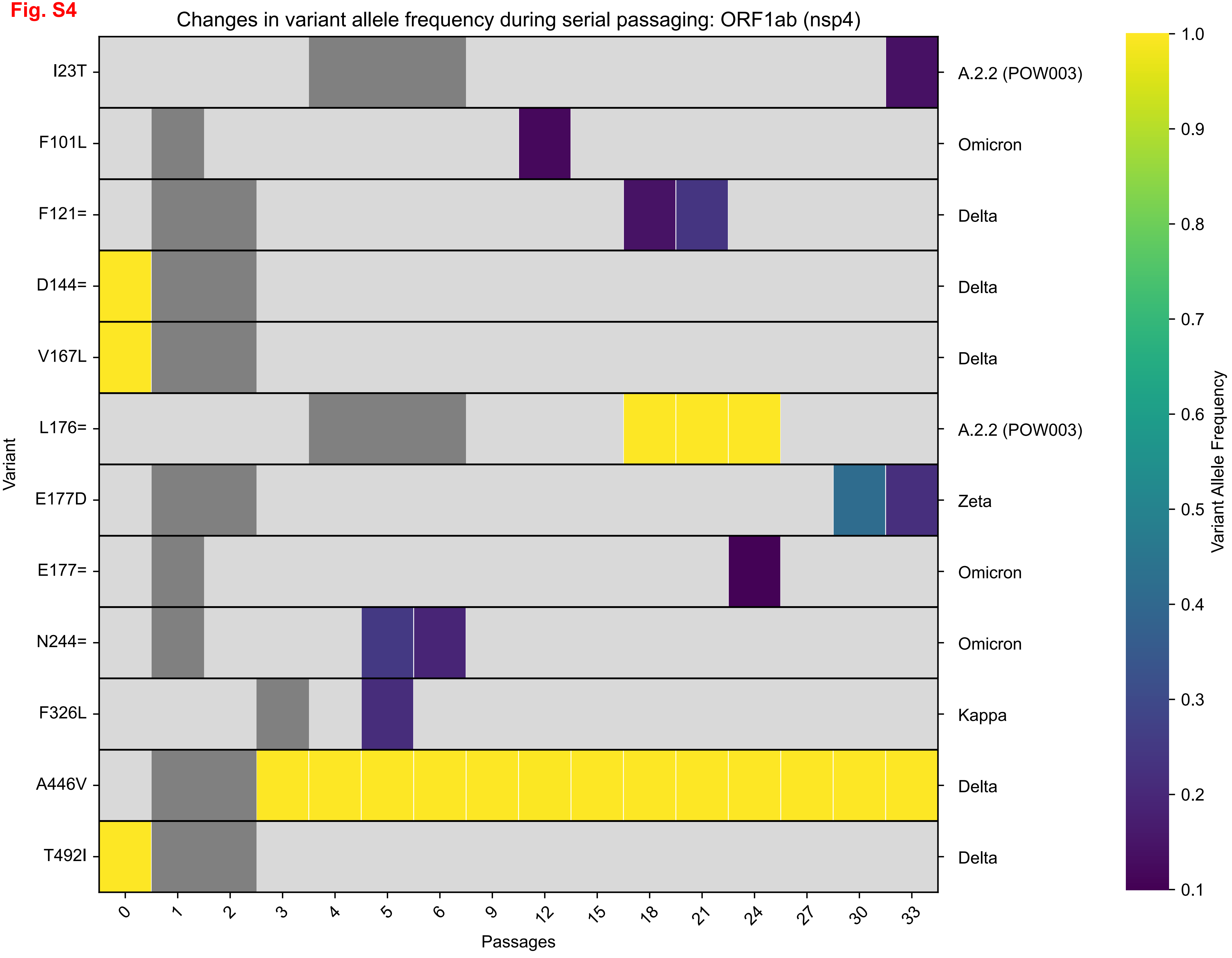

Changes in variant allele frequency during serial passaging: ORF1ab (nsp4)
Variant
I23T
F101L
F121=
D144=
V167L
L176=
E177D
E177=
N244=
F326L
A446V
T492I
A.2.2 (POW003)
Omicron
Delta
Delta
Delta
A.2.2 (POW003)
Zeta
Omicron
Omicron
Kappa
Delta
Delta
0
1
2
3
4
5
6
9
12
15
18
21
24
27
30
33
Passages
Variant Allele Frequency
1.0
0.9
0.8
0.7
0.6
0.5
0.4
0.3
0.2
0.1

**Fig. S5**

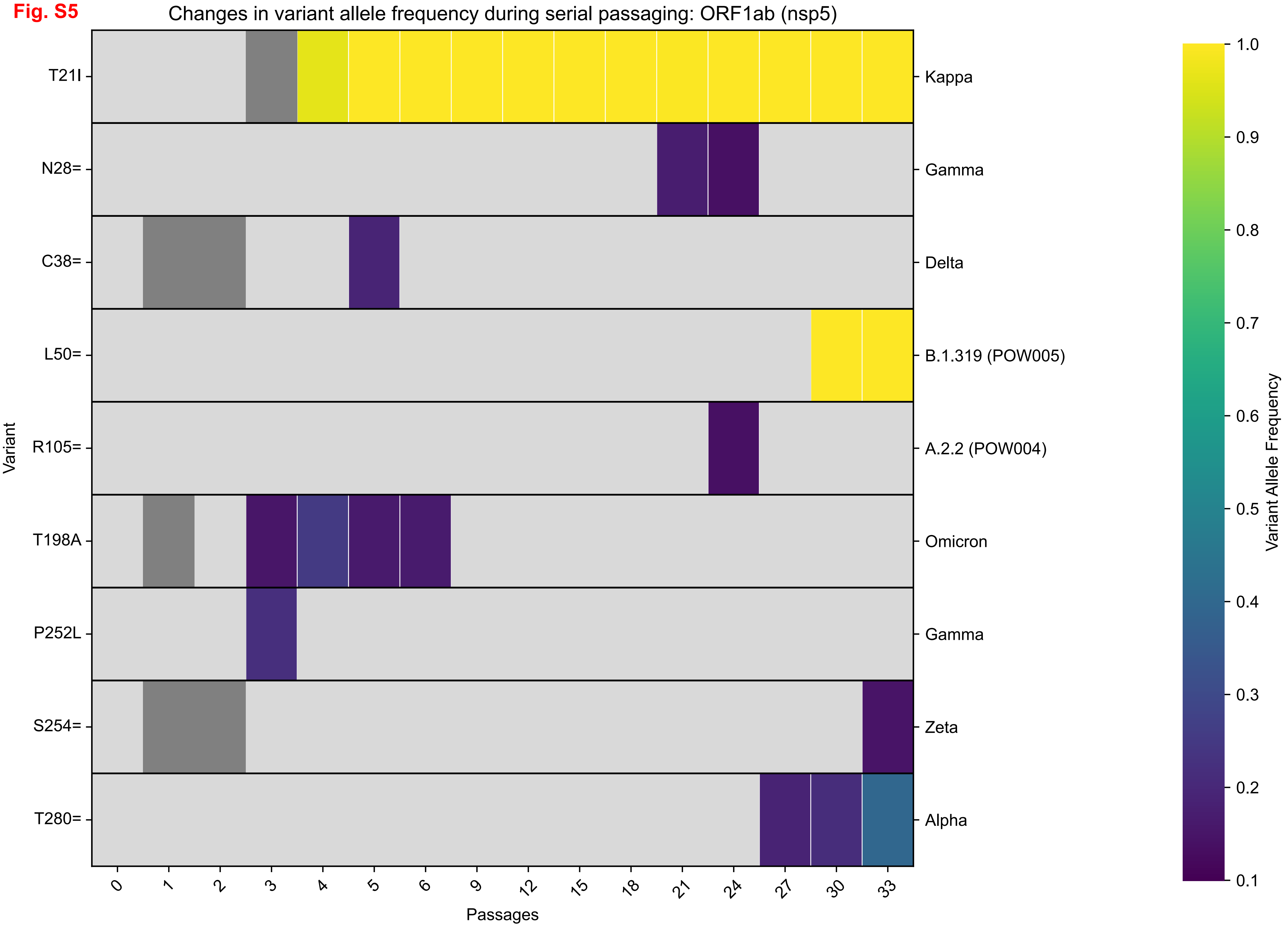
Changes in variant allele frequency during serial passaging: ORF1ab (nsp5)
Variant
T21I
N28=
C38=
L50=
R105=
T198A
P252L
S254=
T280=
Kappa
Gamma
Delta
B.1.319 (POW005)
A.2.2 (POW004)
Omicron
Gamma
Zeta
Alpha
Passages
0
1
2
3
4
5
6
9
12
15
18
21
24
27
30
33
Variant Allele Frequency
1.0
0.9
0.8
0.7
0.6
0.5
0.4
0.3
0.2
0.1

**Fig. S6**

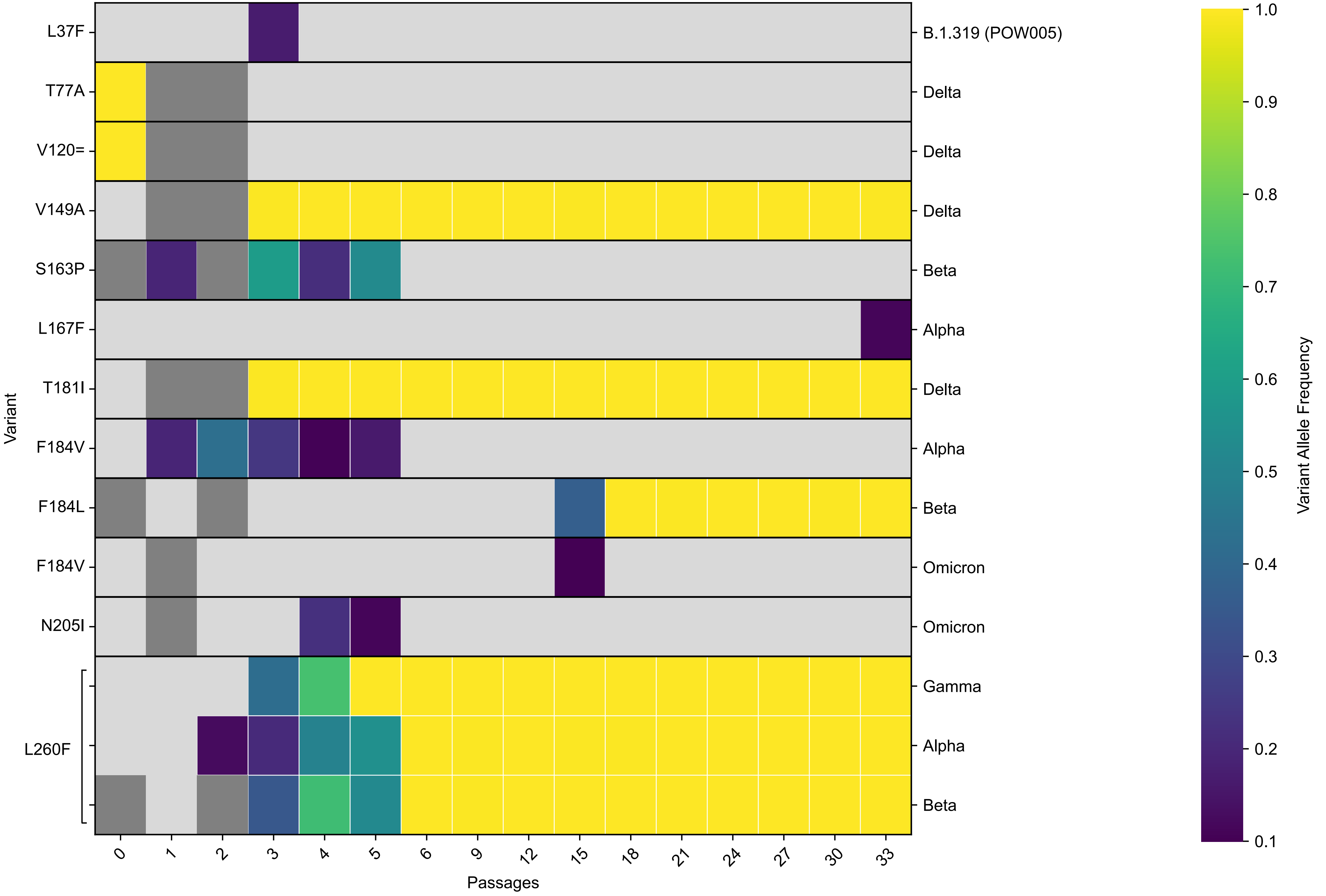
Changes in variant allele frequency during serial passaging: ORF1ab (nsp6)
Variant
L37F
T77A
V120=
V149A
S163P
L167F
T181I
F184V
F184L
F184V
N205I
L260F
B.1.319 (POW005)
Delta
Delta
Delta
Beta
Alpha
Delta
Alpha
Beta
Omicron
Omicron
Gamma
Alpha
Beta
0
1
2
3
4
5
6
9
12
15
18
21
24
27
30
33
Passages
Variant Allele Frequency
1.0
0.9
0.8
0.7
0.6
0.5
0.4
0.3
0.2
0.1

**Fig. S7**

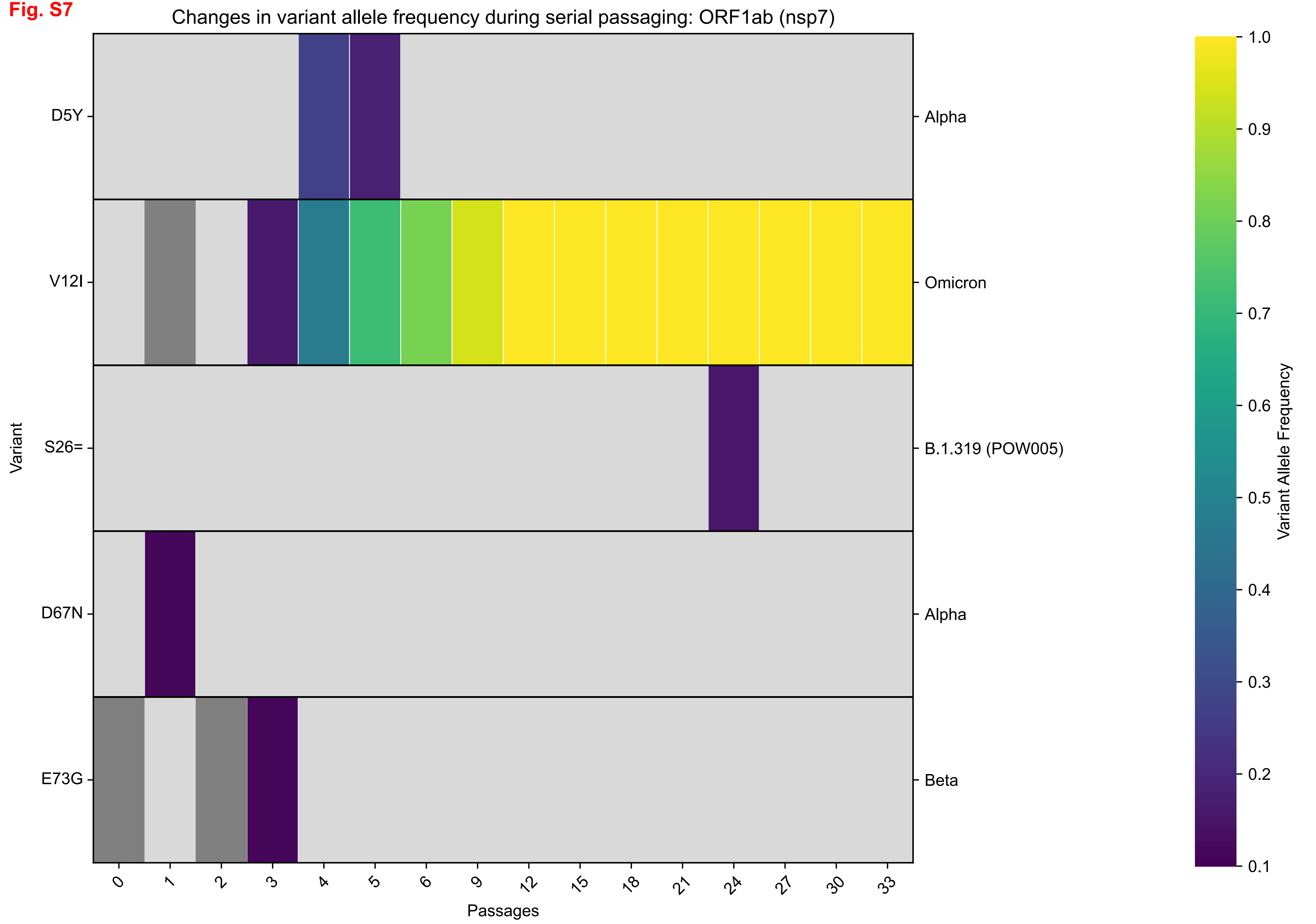

Changes in variant allele frequency during serial passaging: ORF1ab (nsp7)
Variant
D5Y
V12I
S26=
D67N
E73G
Alpha
Omicron
B.1.319 (POW005)
Alpha
Beta
Passages
0
1
2
3
4
5
6
9
12
15
18
21
24
27
30
33
Variant Allele Frequency
1.0
0.9
0.8
0.7
0.6
0.5
0.4
0.3
0.2
0.1

Fig. S8

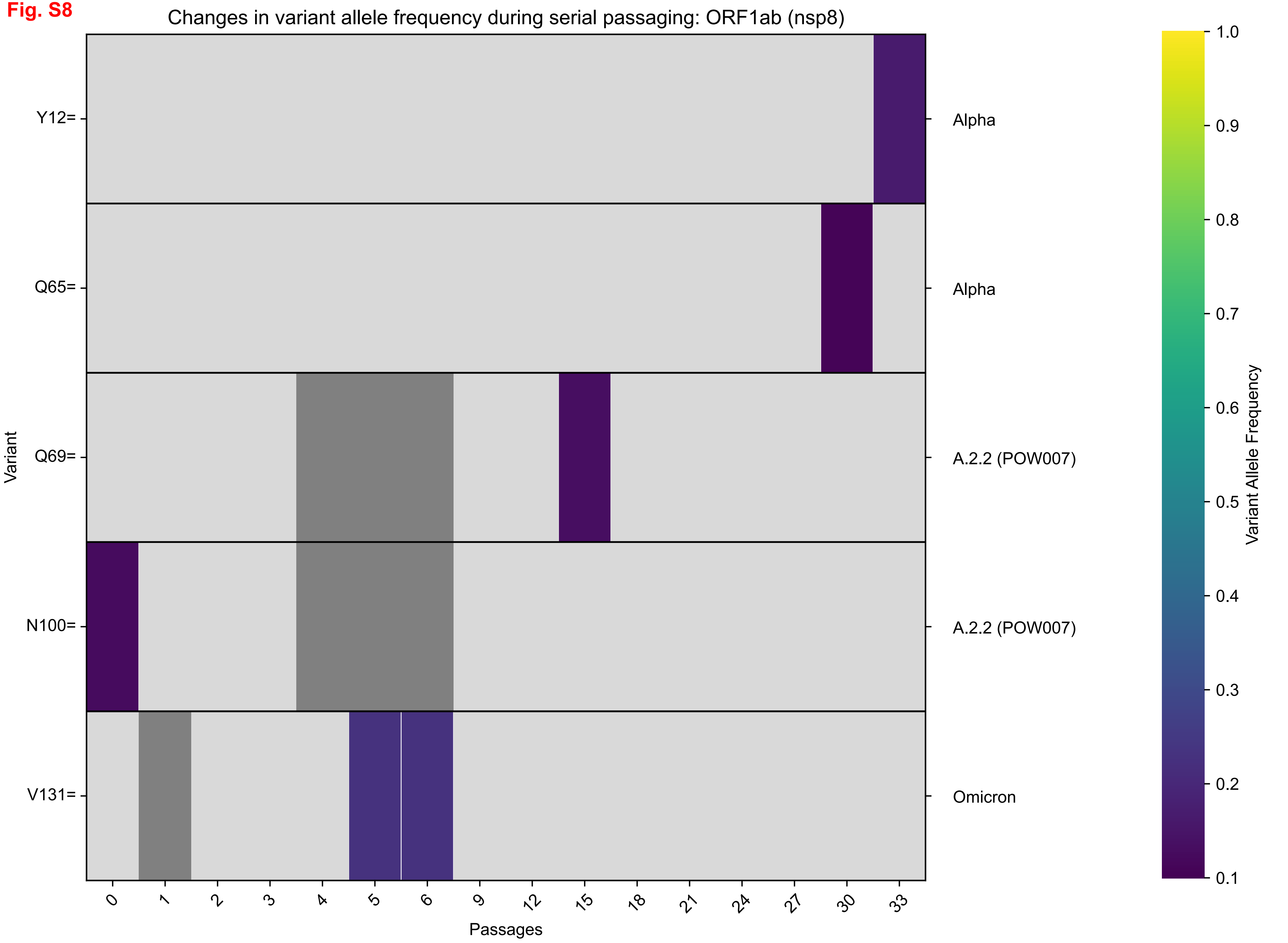
Changes in variant allele frequency during serial passaging: ORF1ab (nsp8)
Variant
Y12=
Q65=
Q69=
N100=
V131=
Alpha
Alpha
A.2.2 (POW007)
A.2.2 (POW007)
Omicron
0
1
2
3
4
5
6
9
12
15
18
21
24
27
30
33
Passages
Variant Allele Frequency
1.0
0.9
0.8
0.7
0.6
0.5
0.4
0.3
0.2
0.1

**Fig. S9**

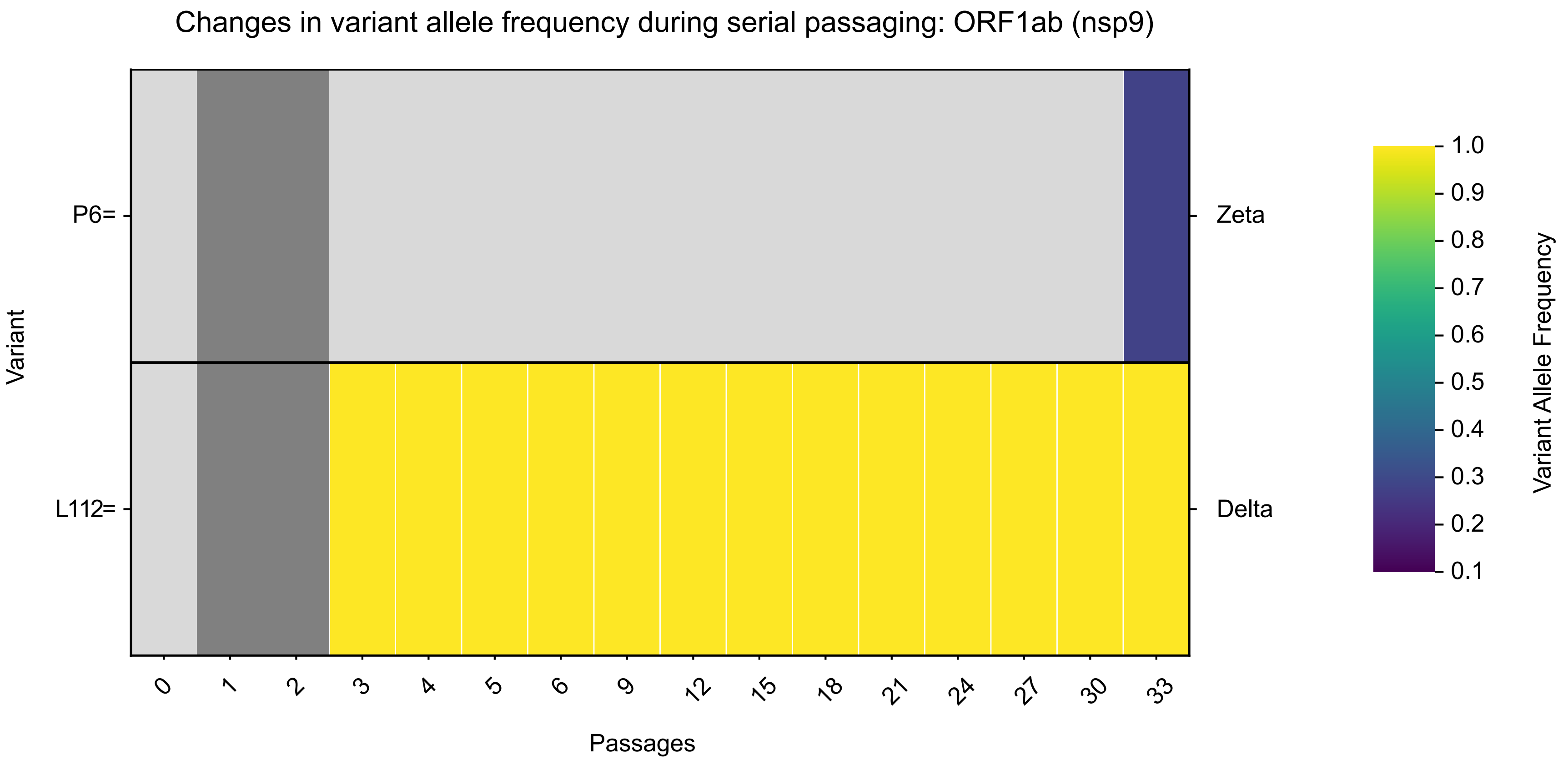

Changes in variant allele frequency during serial passaging: ORF1ab (nsp9)
Variant
P6=
L112=
Zeta
Delta
0
1
2
3
4
5
6
9
12
15
18
21
24
27
30
33
Passages
1.0
0.9
0.8
0.7
0.6
0.5
0.4
0.3
0.2
0.1
Variant Allele Frequency

**Fig. S10**

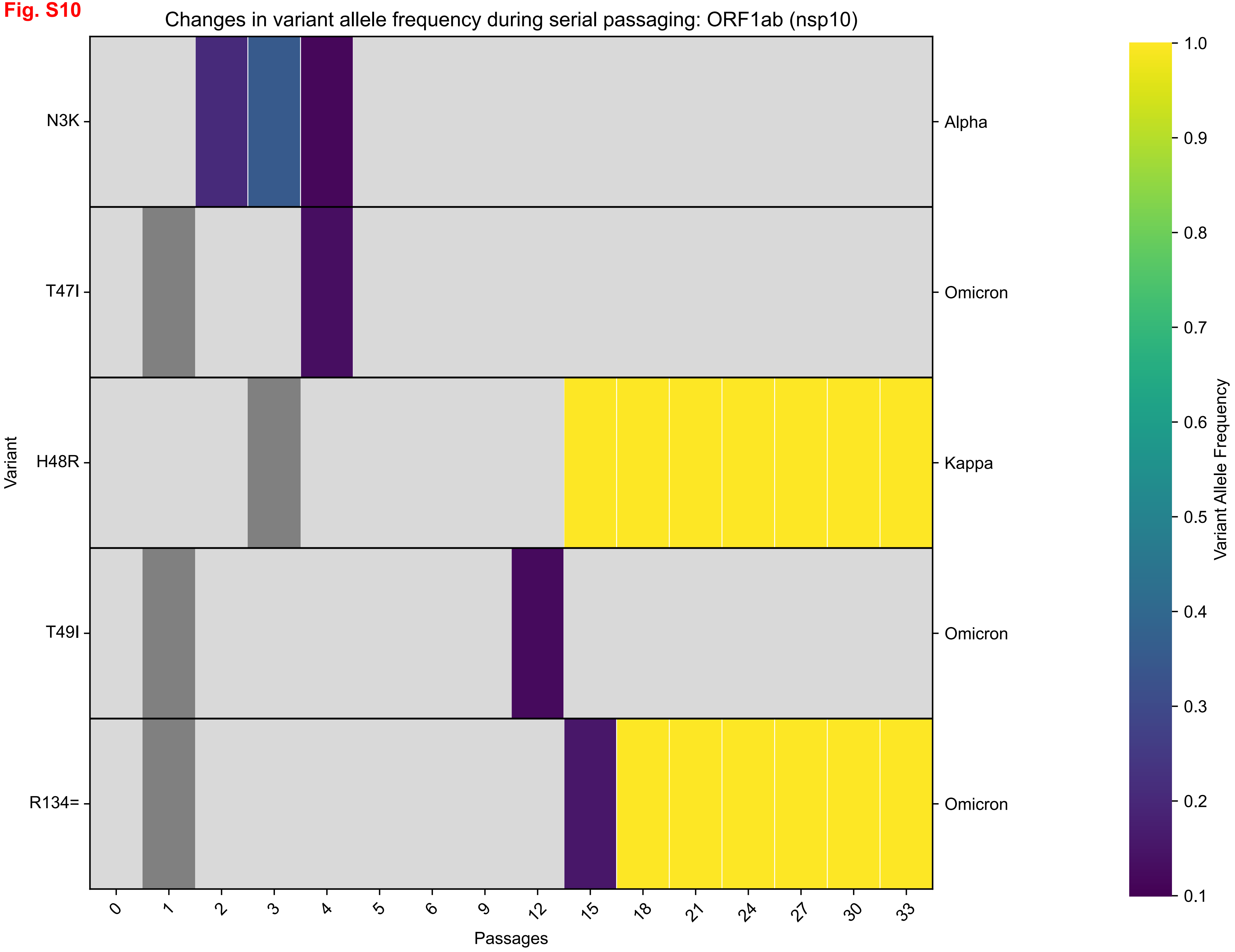

Changes in variant allele frequency during serial passaging: ORF1ab (nsp10)
Variant
N3K
T47I
H48R
T49I
R134=
Alpha
Omicron
Kappa
Omicron
Omicron
Passages
0
1
2
3
4
5
6
9
12
15
18
21
24
27
30
33
Variant Allele Frequency
1.0
0.9
0.8
0.7
0.6
0.5
0.4
0.3
0.2
0.1

**Fig. S11**

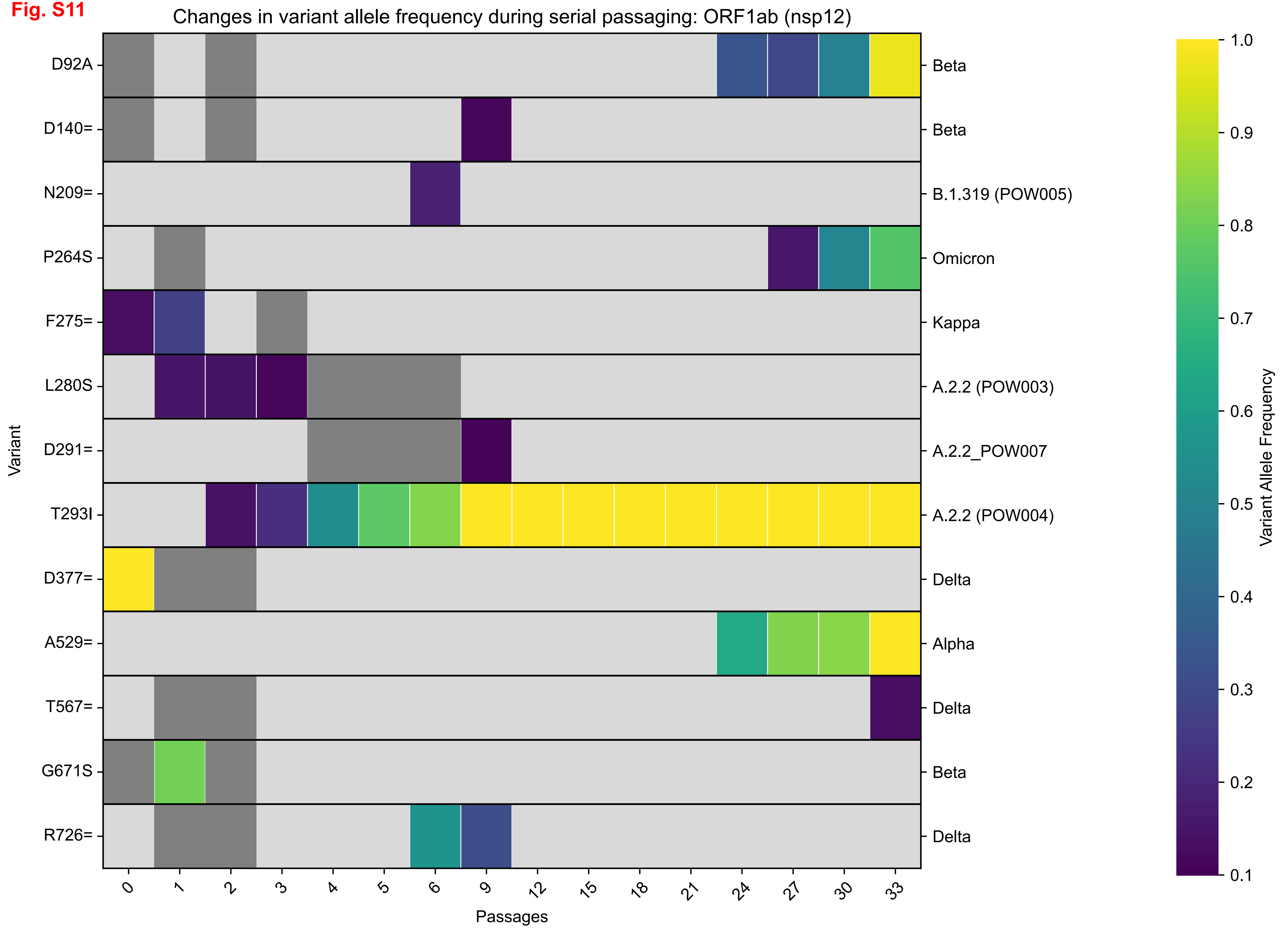
Changes in variant allele frequency during serial passaging: ORF1ab (nsp12)
Variant
D92A
D140=
N209=
P264S
F275=
L280S
D291=
T293I
D377=
A529=
T567=
G671S
R726=
Beta
Beta
B.1.319 (POW005)
Omicron
Kappa
A.2.2 (POW003)
A.2.2_POW007
A.2.2 (POW004)
Delta
Alpha
Delta
Beta
Delta
0
1
2
3
4
5
6
9
12
15
18
21
24
27
30
33
Passages
Variant Allele Frequency
1.0
0.9
0.8
0.7
0.6
0.5
0.4
0.3
0.2
0.1

Fig. S12

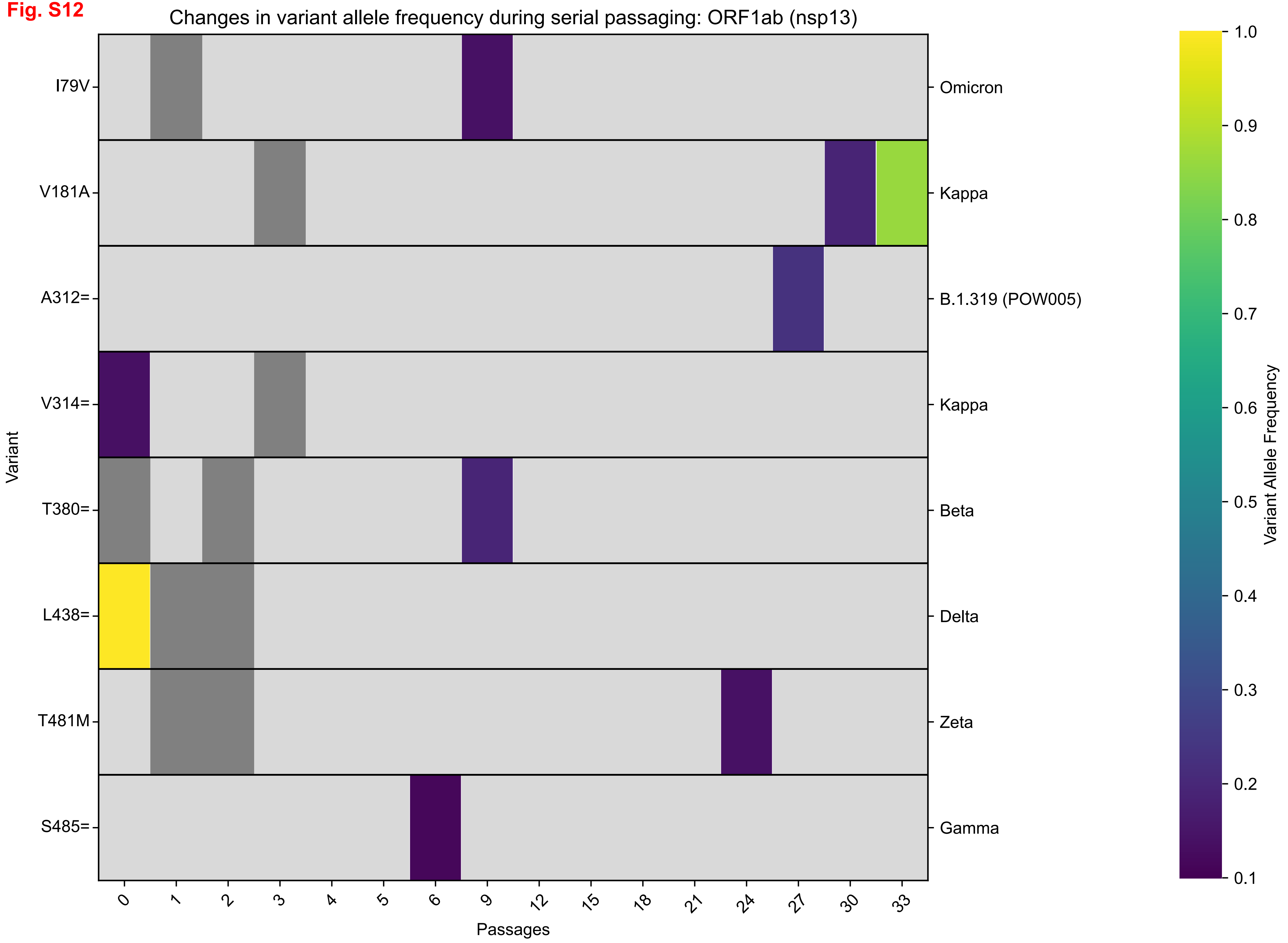
Changes in variant allele frequency during serial passaging: ORF1ab (nsp13)
I79V
V181A
A312=
V314=
T380=
L438=
T481M
S485=
Variant
Omicron
Kappa
B.1.319 (POW005)
Kappa
Beta
Delta
Zeta
Gamma
0
1
2
3
4
5
6
9
12
15
18
21
24
27
30
33
Passages
1.0
0.9
0.8
0.7
0.6
0.5
0.4
0.3
0.2
0.1
Variant Allele Frequency

**Fig. S13**

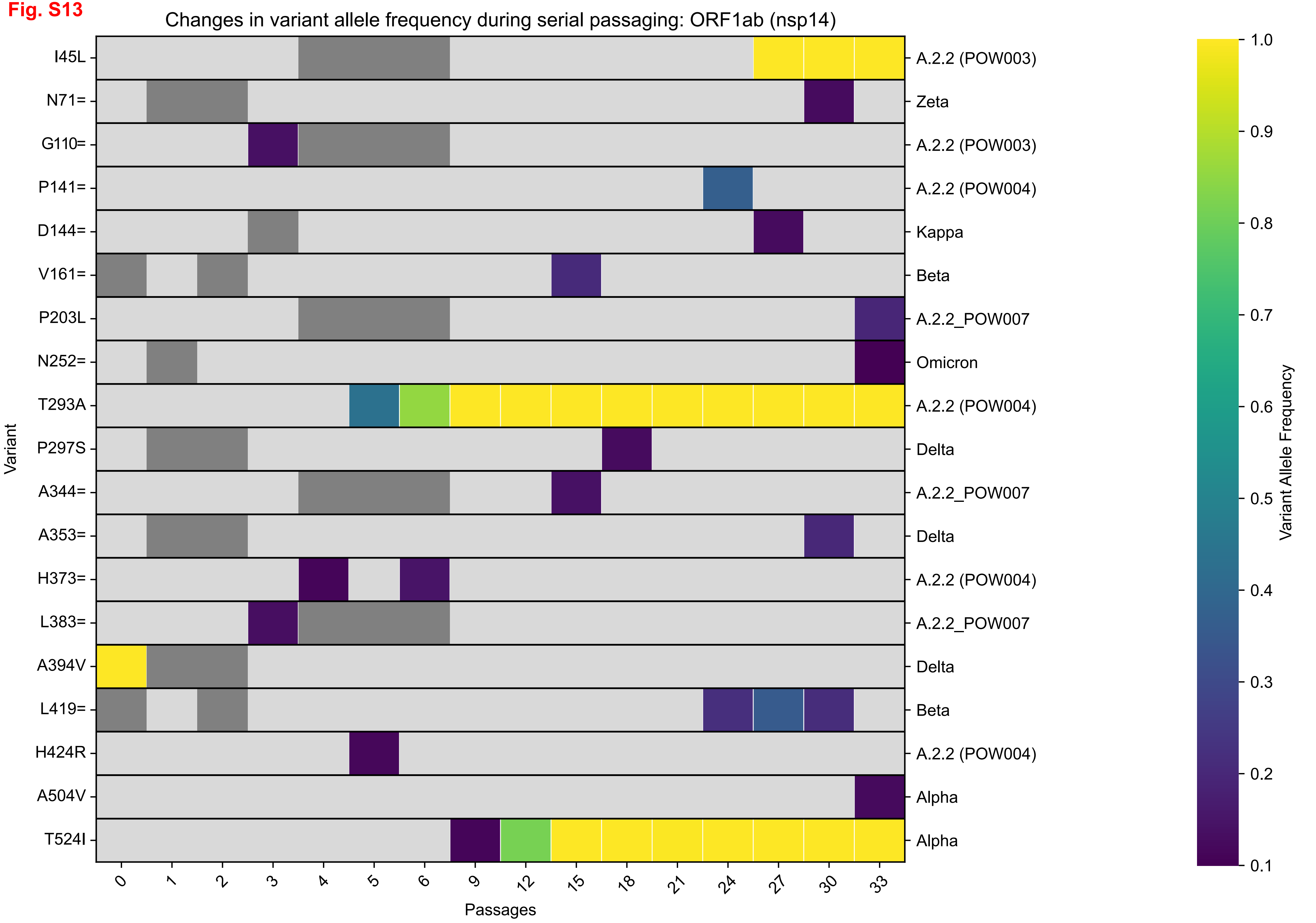
Changes in variant allele frequency during serial passaging: ORF1ab (nsp14)
Variant
I45L
N71=
G110=
P141=
D144=
V161=
P203L
N252=
T293A
P297S
A344=
A353=
H373=
L383=
A394V
L419=
H424R
A504V
T524I
A.2.2 (POW003)
Zeta
A.2.2 (POW003)
A.2.2 (POW004)
Kappa
Beta
A.2.2_POW007
Omicron
A.2.2 (POW004)
Delta
A.2.2_POW007
Delta
A.2.2 (POW004)
A.2.2_POW007
Delta
Beta
A.2.2 (POW004)
Alpha
Alpha
0
1
2
3
4
5
6
9
12
15
18
21
24
27
30
33
Passages
Variant Allele Frequency
1.0
0.9
0.8
0.7
0.6
0.5
0.4
0.3
0.2
0.1

Fig. S14

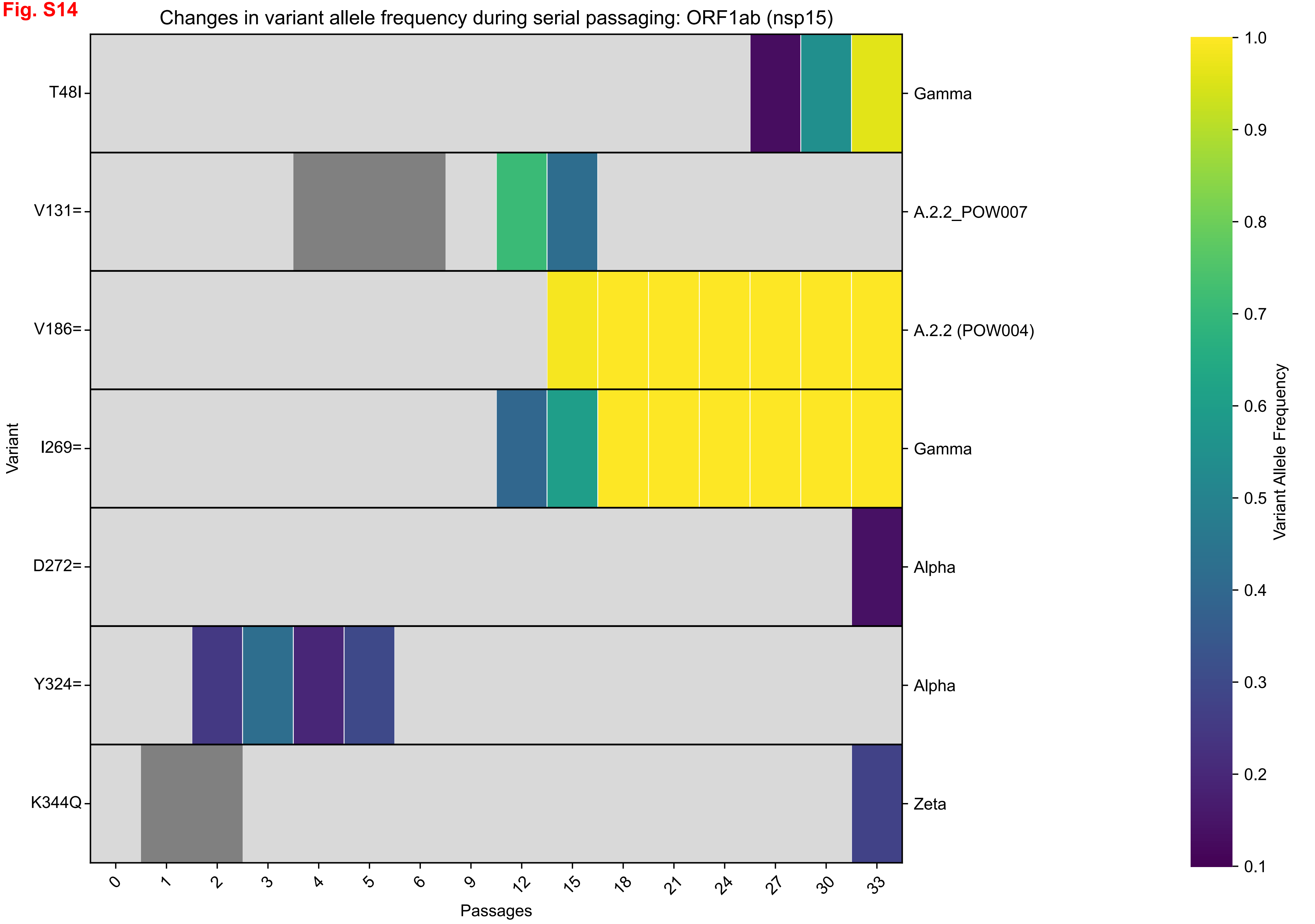

Changes in variant allele frequency during serial passaging: ORF1ab (nsp15)
Variant
T48I
V131=
V186=
I269=
D272=
Y324=
K344Q
Gamma
A.2.2_POW007
A.2.2 (POW004)
Gamma
Alpha
Alpha
Zeta
0
1
2
3
4
5
6
9
12
15
18
21
24
27
30
33
Passages
Variant Allele Frequency
1.0
0.9
0.8
0.7
0.6
0.5
0.4
0.3
0.2
0.1

**Fig. S15**

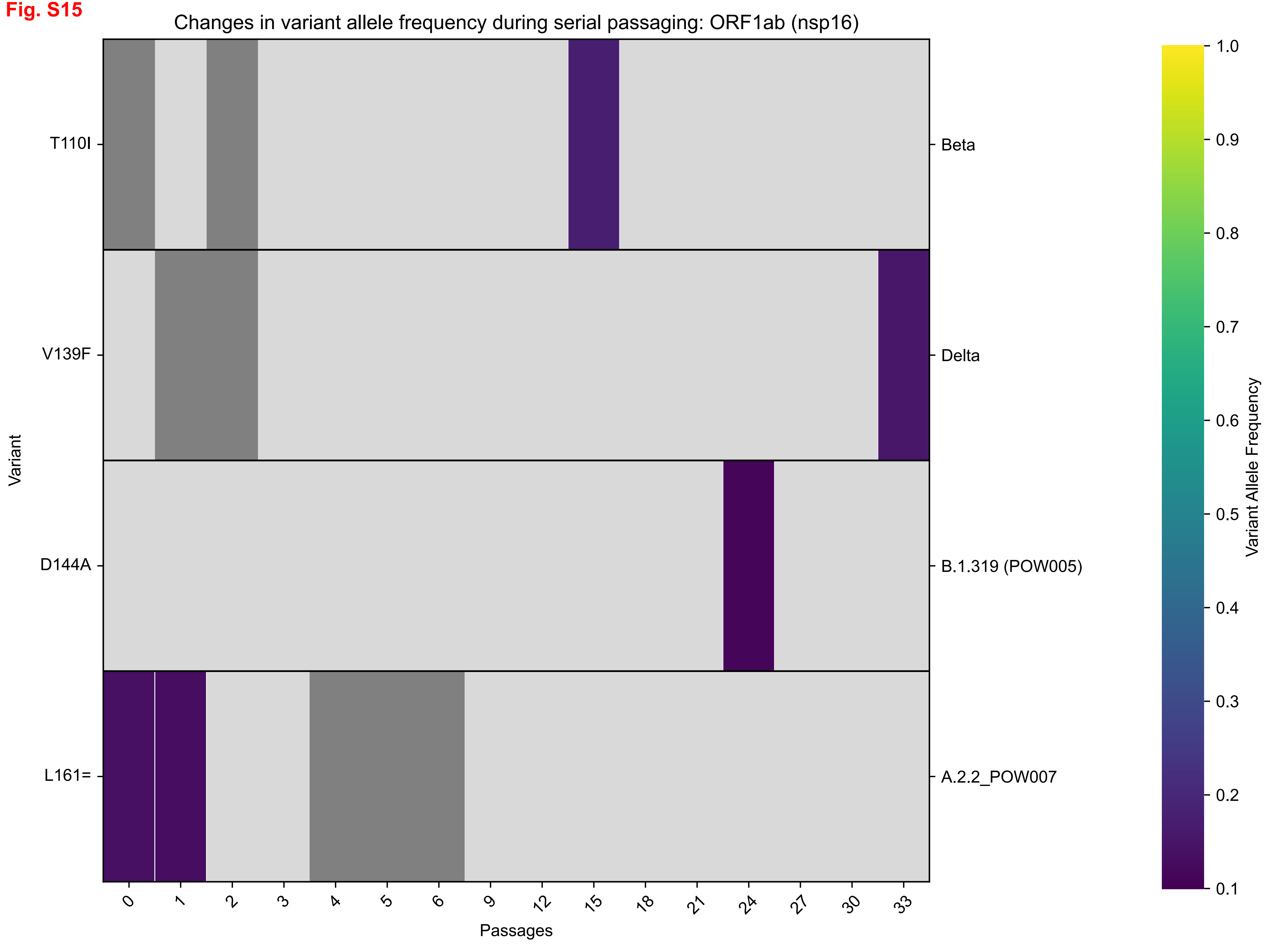
Changes in variant allele frequency during serial passaging: ORF1ab (nsp16)
T110I
V139F
D144A
L161=
Variant
Beta
Delta
B.1.319 (POW005)
A.2.2_POW007
0
1
2
3
4
5
6
9
12
15
18
21
24
27
30
33
Passages
1.0
0.9
0.8
0.7
0.6
0.5
0.4
0.3
0.2
0.1
Variant Allele Frequency

Fig. S16

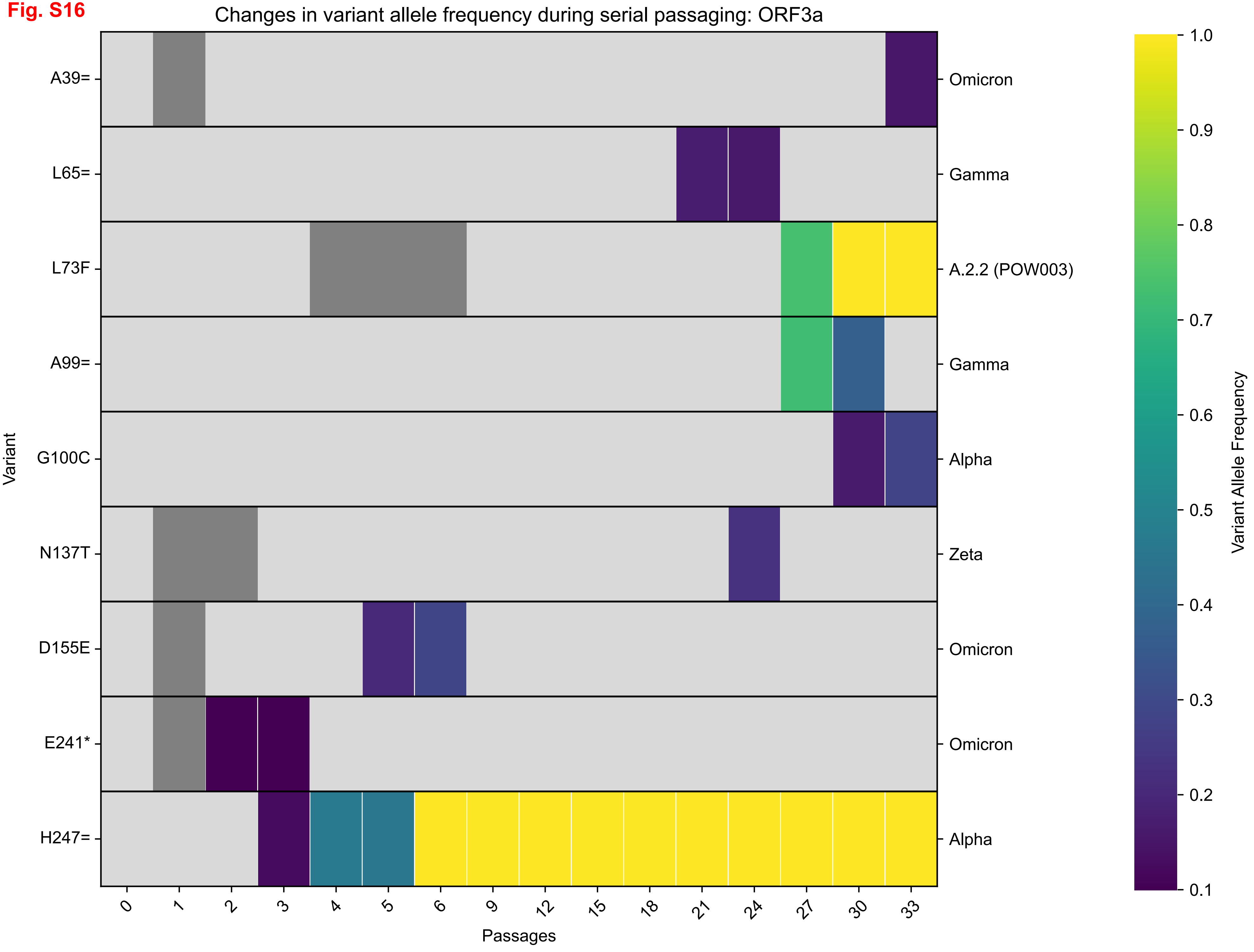
Changes in variant allele frequency during serial passaging: ORF3a
A39=
L65=
L73F
A99=
G100C
N137T
D155E
E241*
H247=
Omicron
Gamma
A.2.2 (POW003)
Gamma
Alpha
Zeta
Omicron
Omicron
Alpha
Variant
0
1
2
3
4
5
6
9
12
15
18
21
24
27
30
33
Passages
1.0
0.9
0.8
0.7
0.6
0.5
0.4
0.3
0.2
0.1
Variant Allele Frequency

**Fig. S17**

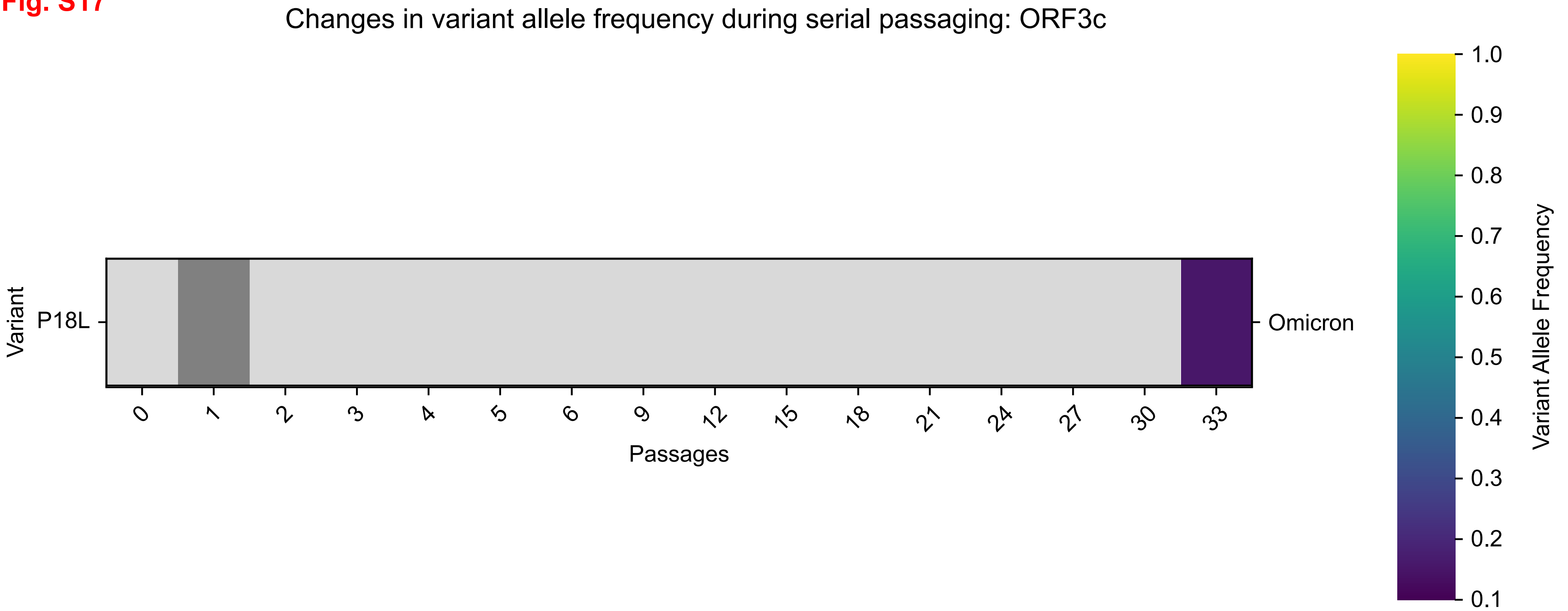
Changes in variant allele frequency during serial passaging: ORF3c
Variant
P18L
Omicron
0
1
2
3
4
5
6
9
12
15
18
21
24
27
30
33
Passages
1.0
0.9
0.8
0.7
0.6
0.5
0.4
0.3
0.2
0.1
Variant Allele Frequency

**Fig. S18**

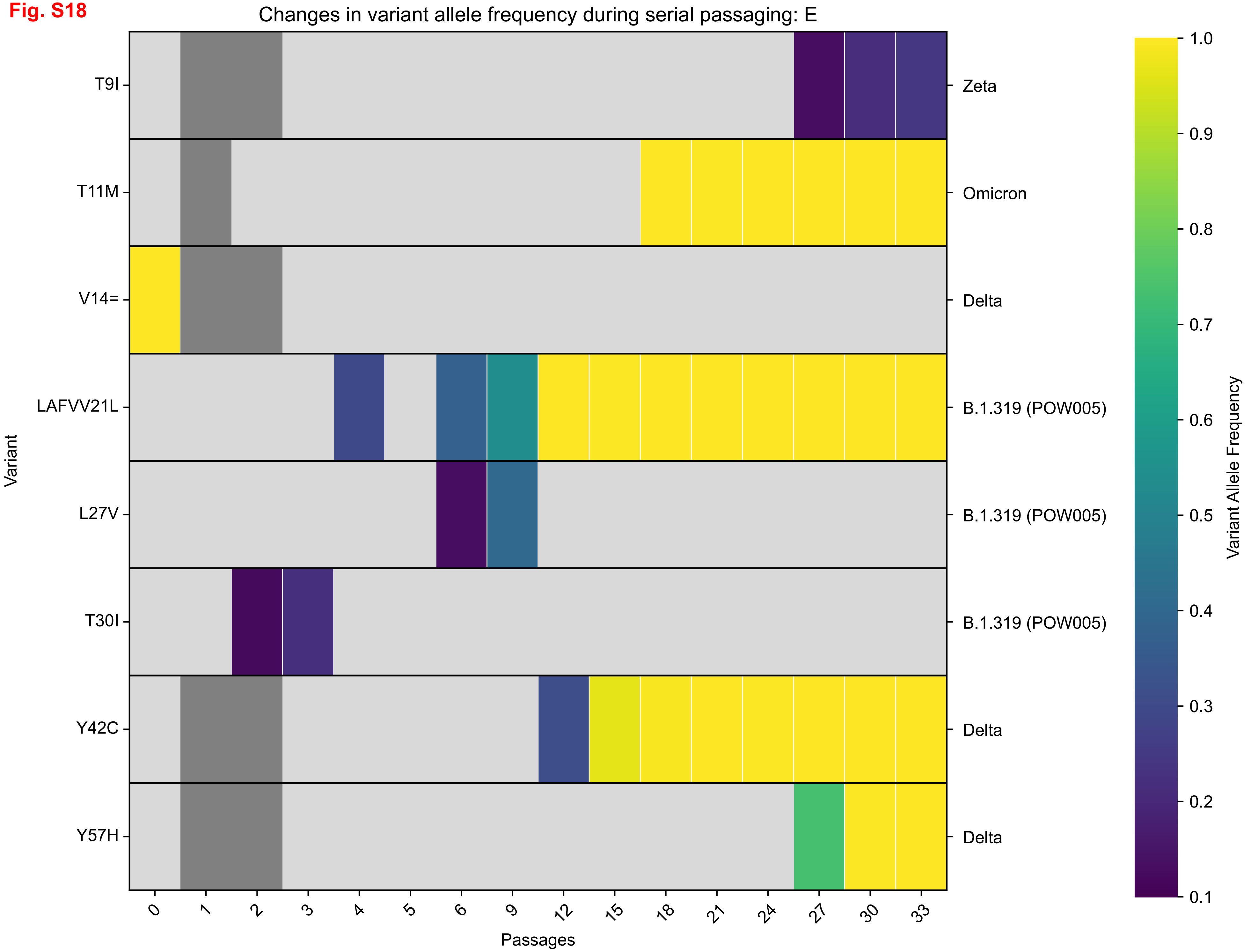
Changes in variant allele frequency during serial passaging: E
Variant
T9I
T11M
V14=
LAFVV21L
L27V
T30I
Y42C
Y57H
Zeta
Omicron
Delta
B.1.319 (POW005)
B.1.319 (POW005)
B.1.319 (POW005)
Delta
Delta
Passages
0
1
2
3
4
5
6
9
12
15
18
21
24
27
30
33
Variant Allele Frequency
1.0
0.9
0.8
0.7
0.6
0.5
0.4
0.3
0.2
0.1

Fig. S19

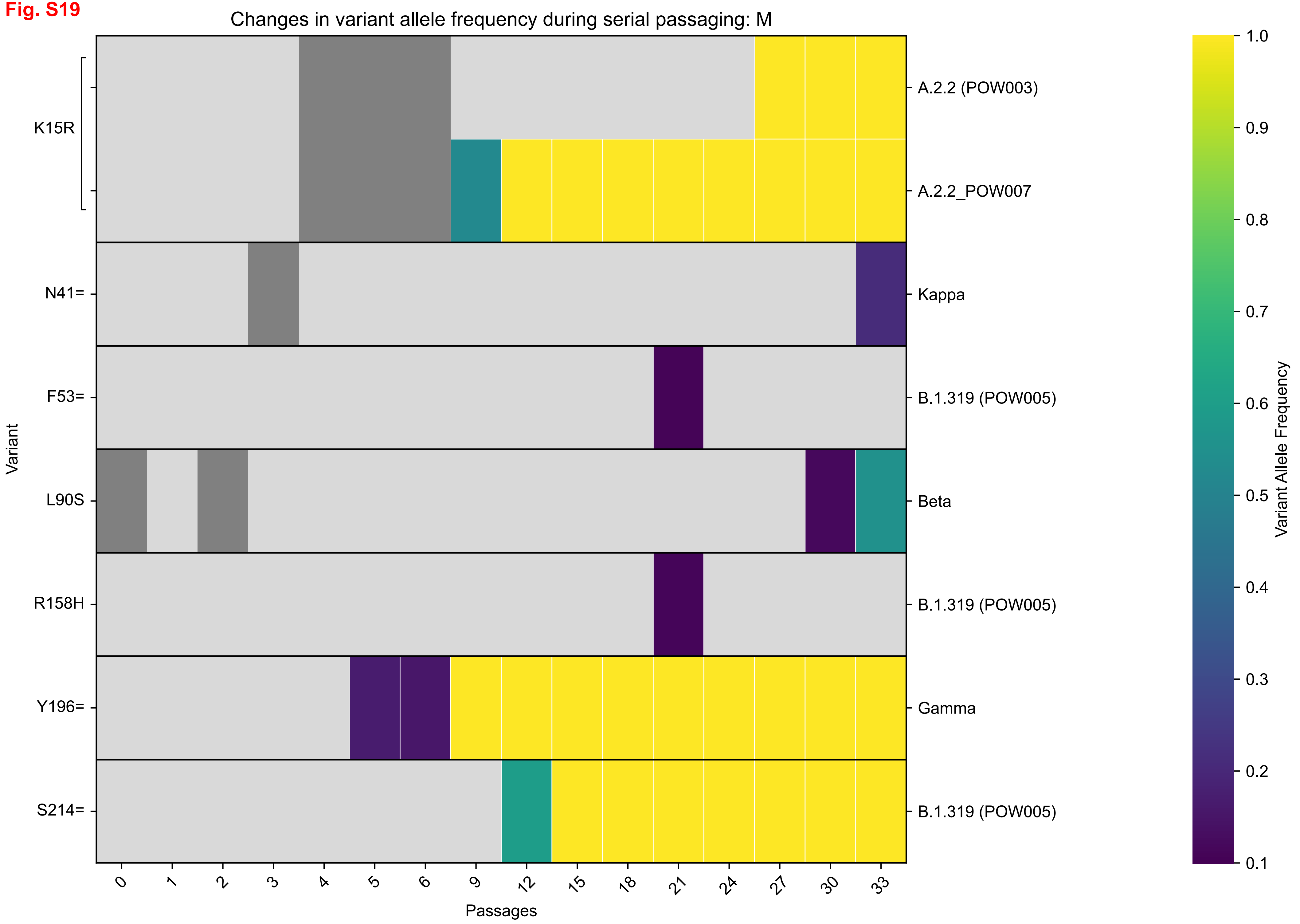
Changes in variant allele frequency during serial passaging: M
K15R
N41=
F53=
L90S
R158H
Y196=
S214=
Variant
A.2.2 (POW003)
A.2.2_POW007
Kappa
B.1.319 (POW005)
Beta
B.1.319 (POW005)
Gamma
B.1.319 (POW005)
0
1
2
3
4
5
6
9
12
15
18
21
24
27
30
33
Passages
1.0
0.9
0.8
0.7
0.6
0.5
0.4
0.3
0.2
0.1
Variant Allele Frequency

Fig. S20

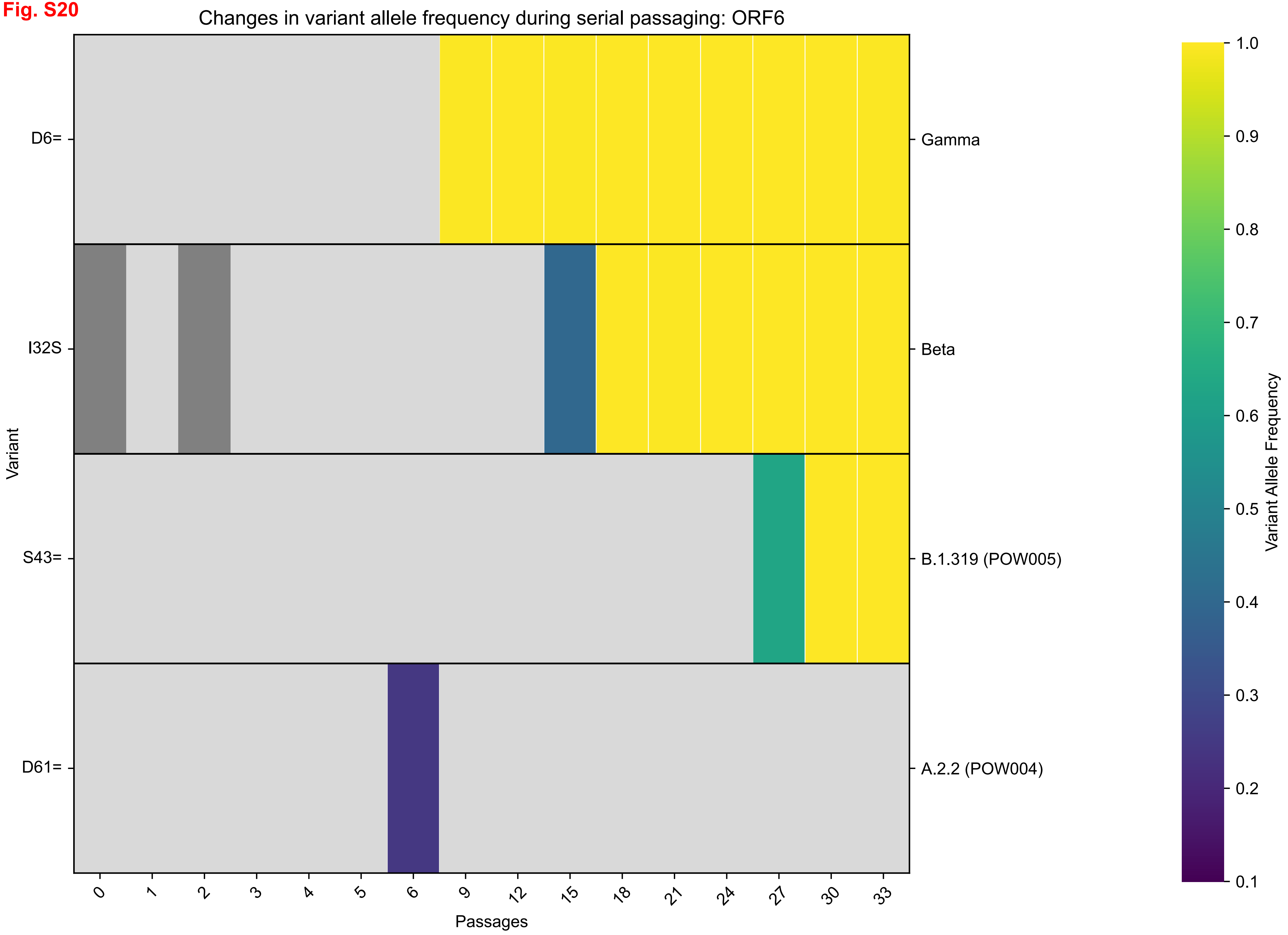
Changes in variant allele frequency during serial passaging: ORF6
Variant
D6=
I32S
S43=
D61=
Gamma
Beta
B.1.319 (POW005)
A.2.2 (POW004)
Passages
0
1
2
3
4
5
6
9
12
15
18
21
24
27
30
33
Variant Allele Frequency
1.0
0.9
0.8
0.7
0.6
0.5
0.4
0.3
0.2
0.1

**Fig. S21**

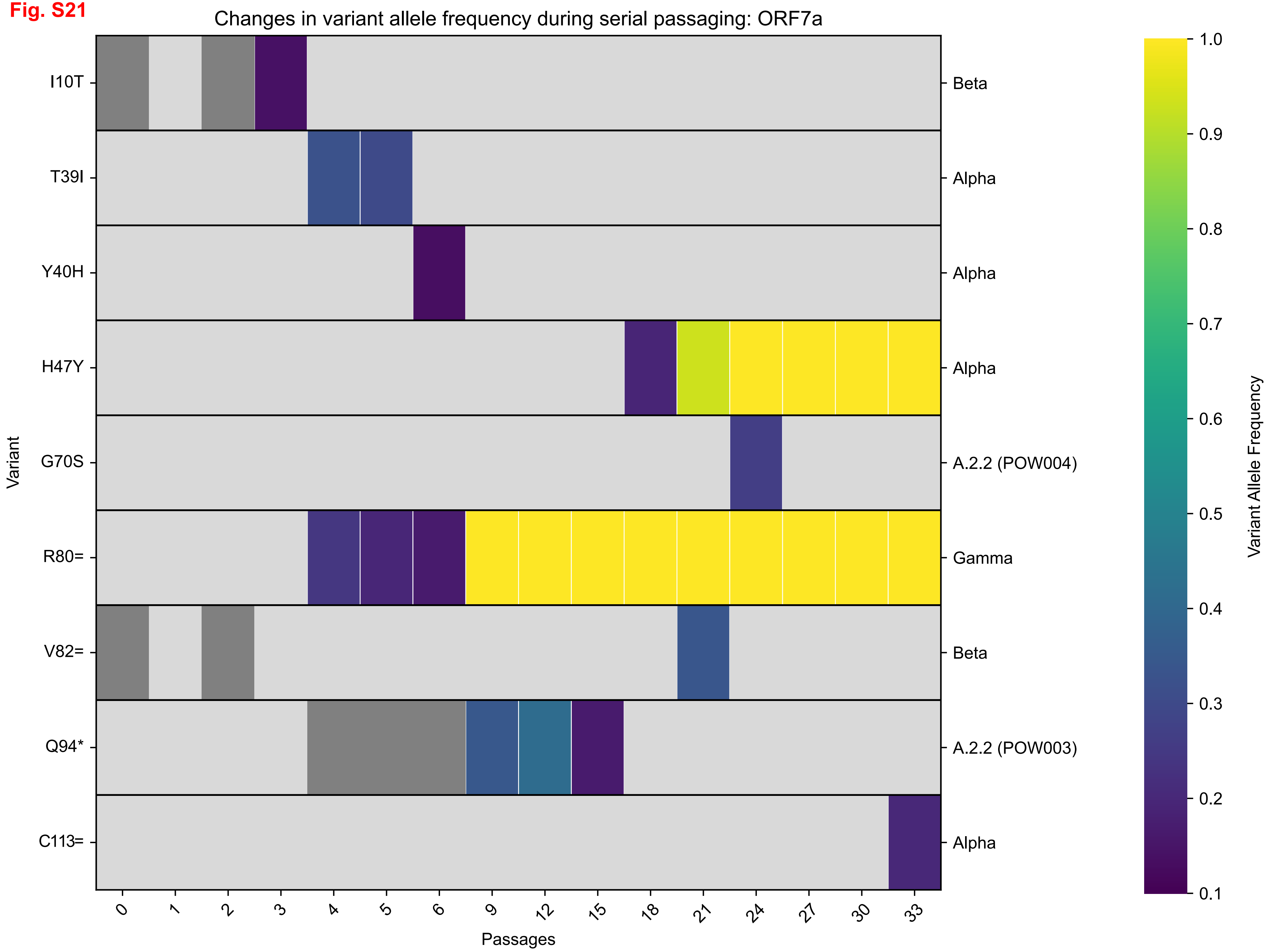

Changes in variant allele frequency during serial passaging: ORF7a
Variant
I10T
T39I
Y40H
H47Y
G70S
R80=
V82=
Q94*
C113=
Beta
Alpha
Alpha
Alpha
A.2.2 (POW004)
Gamma
Beta
A.2.2 (POW003)
Alpha
0
1
2
3
4
5
6
9
12
15
18
21
24
27
30
33
Passages
Variant Allele Frequency
1.0
0.9
0.8
0.7
0.6
0.5
0.4
0.3
0.2
0.1

**Fig. S22**

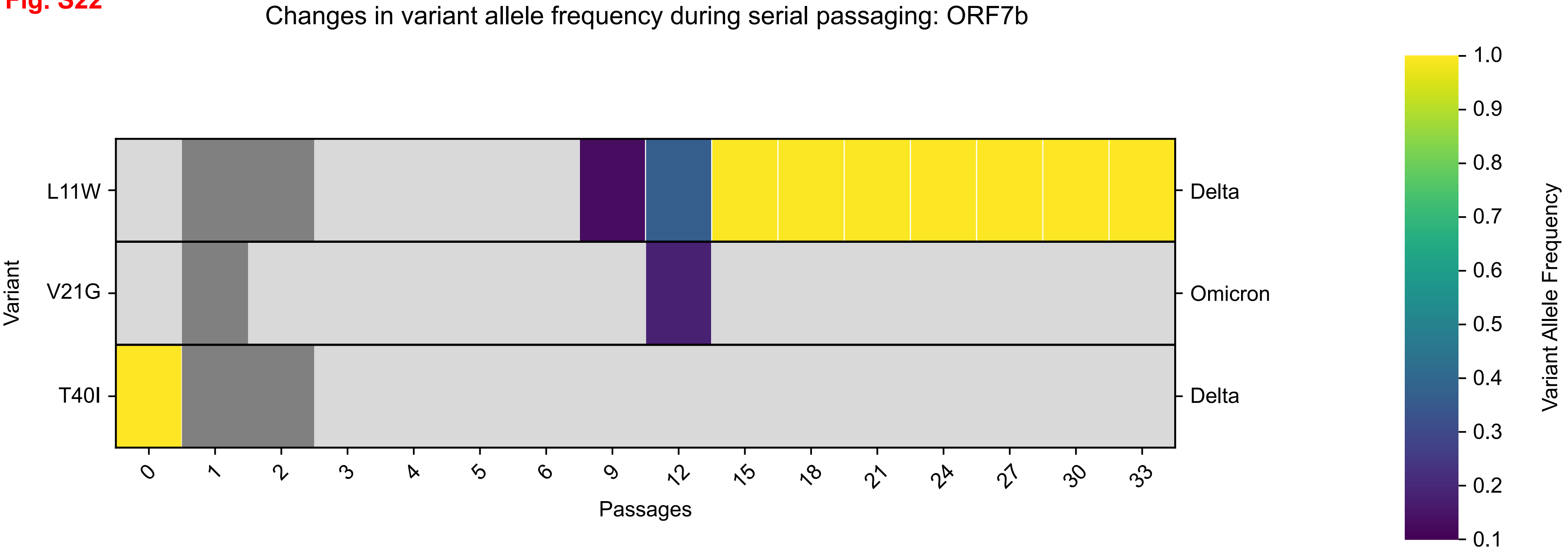

Changes in variant allele frequency during serial passaging: ORF7b
L11W
V21G
T40I
Variant
Delta
Omicron
Delta
0
1
2
3
4
5
6
9
12
15
18
21
24
27
30
33
Passages
1.0
0.9
0.8
0.7
0.6
0.5
0.4
0.3
0.2
0.1
Variant Allele Frequency

**Fig. S23**

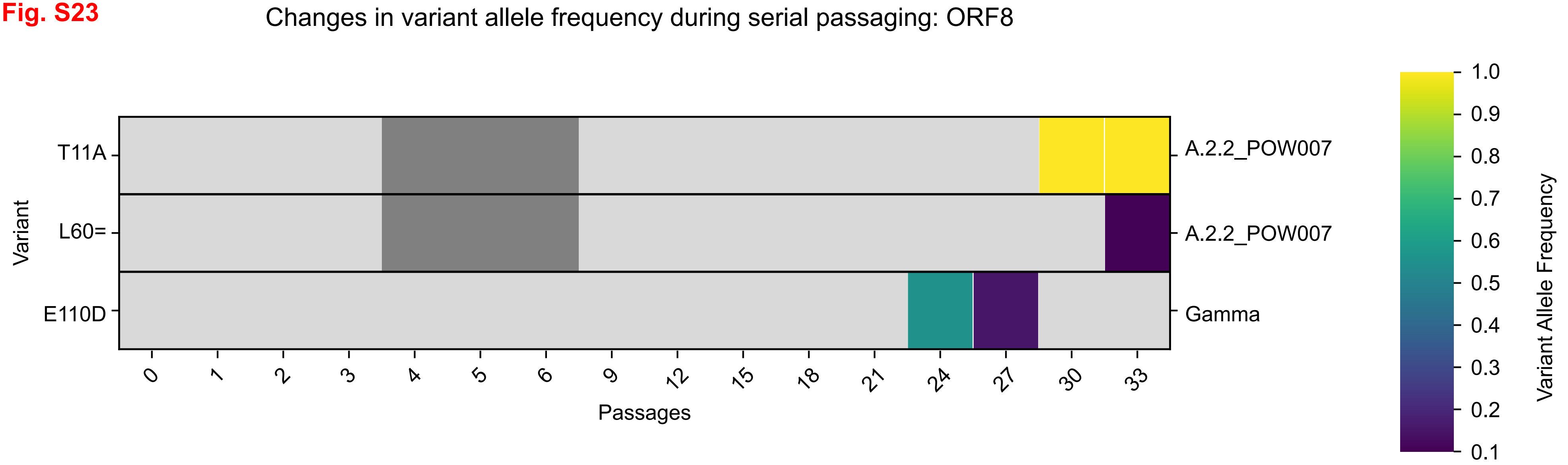
Changes in variant allele frequency during serial passaging: ORF8
Variant
T11A
L60=
E110D
A.2.2_POW007
A.2.2_POW007
Gamma
0
1
2
3
4
5
6
9
12
15
18
21
24
27
30
33
Passages
1.0
0.9
0.8
0.7
0.6
0.5
0.4
0.3
0.2
0.1
Variant Allele Frequency

**Fig. S24**

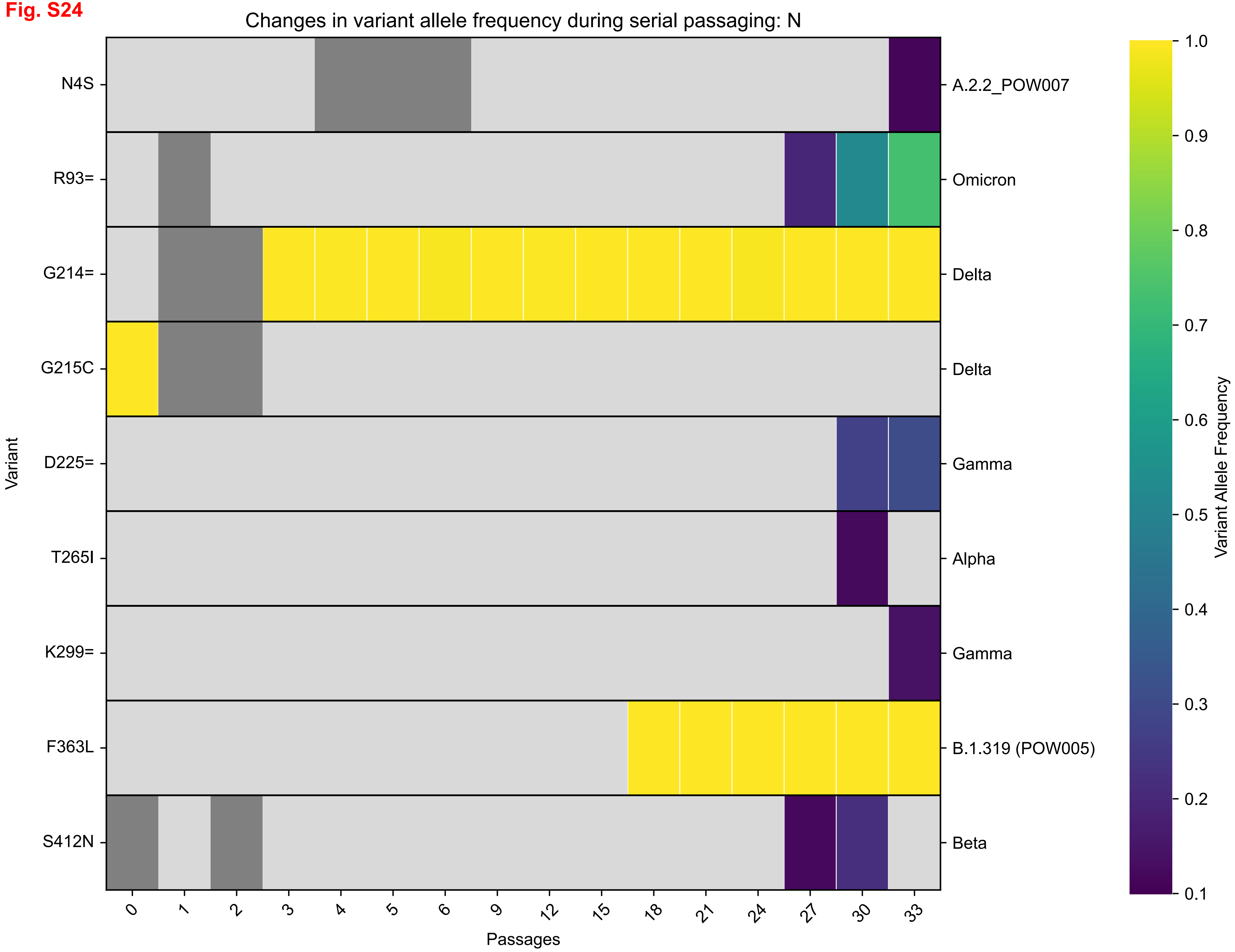

Changes in variant allele frequency during serial passaging: N
Variant
N4S
R93=
G214=
G215C
D225=
T265I
K299=
F363L
S412N
A.2.2_POW007
Omicron
Delta
Delta
Gamma
Alpha
Gamma
B.1.319 (POW005)
Beta
0
1
2
3
4
5
6
9
12
15
18
21
24
27
30
33
Passages
Variant Allele Frequency
1.0
0.9
0.8
0.7
0.6
0.5
0.4
0.3
0.2
0.1

**Fig. S25**

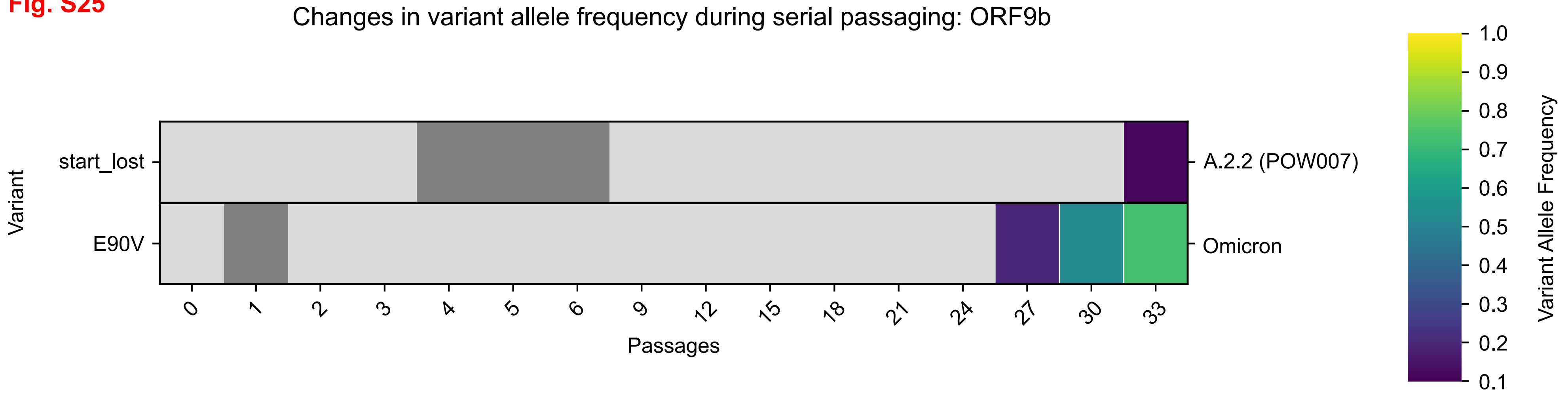
Changes in variant allele frequency during serial passaging: ORF9b
Variant
start_lost
E90V
A.2.2 (POW007)
Omicron
0
1
2
3
4
5
6
9
12
15
18
21
24
27
30
33
Passages
Variant Allele Frequency
1.0
0.9
0.8
0.7
0.6
0.5
0.4
0.3
0.2
0.1

**Fig. S26**

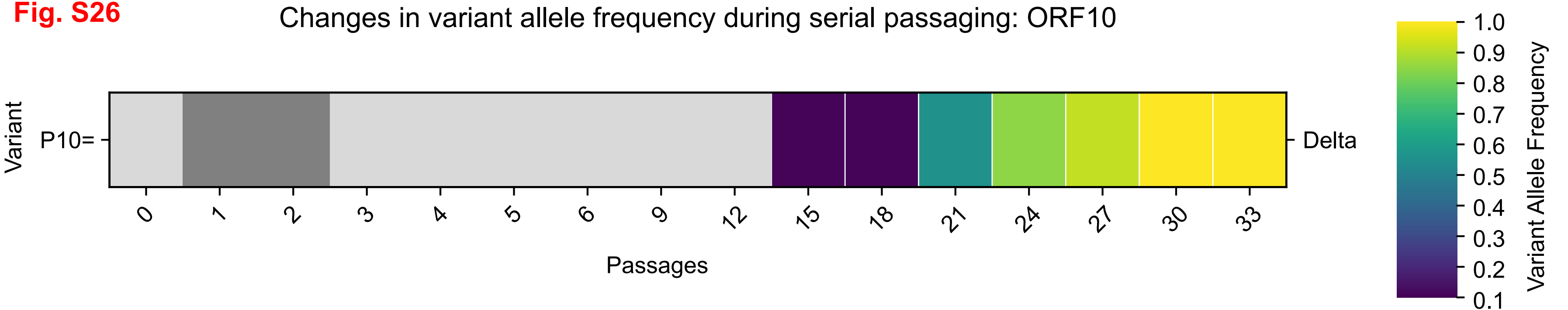
Changes in variant allele frequency during serial passaging: ORF10
Variant
P10=
Delta
0
1
2
3
4
5
6
9
12
15
18
21
24
27
30
33
Passages
1.0
0.9
0.8
0.7
0.6
0.5
0.4
0.3
0.2
0.1
Variant Allele Frequency
