## Supplementary material for "Long-term serial passaging of SARS-CoV-2 reveals signatures of convergent evolution": Supplementary_Captions.docx

**Figures**

Figure S1–26: Heatmaps depicting the variants detected in each gene of SARS-CoV-2 (except spike, which is included in the main text) in each passage line throughout the course of serial passaging. The x-axis refers to the passage number (range: 0 [clinical isolate] to 33). On the primary y-axis (lefthand side) the inferred amino acid consequences of variants are listed (see Supplementary Table S2 for further detail). On the secondary y-axis (righthand side) the data in a given row is linked to the passage line from which it derives. Each ‘cell’ within the heatmap is shaded based on the variant allele frequency of a given variant in a given passage number within that passage line (light grey: undetected). Any cells that are dark grey represent cases where there is no sequencing data for that passage number. Convergence among passage lines is indicated by brackets linking a given variant to multiple passage lines. The title of each supplementary figure details the gene from which the results derive.

**Tables**

Table S1: Quality control and analysis metrics for the sequencing data and consensus genome assemblies for all samples within the study.

Table S2: Full combined results from the ‘vartracker’ pipeline (see main text) for all passage lines within the study.
